# Ozanimod Attenuates Human Cerebrovascular Endothelial Derived MMP-9 Activity and Preserves Barrier Properties Following In Vitro Acute Ischemic Injury

**DOI:** 10.1101/2023.02.01.526738

**Authors:** Trevor S. Wendt, Rayna J. Gonzales

**Affiliations:** Department of Basic Medical Sciences, University of Arizona College of Medicine, Phoenix

**Keywords:** ozanimod, sphingosine-1-phosphate receptor 1, human brain endothelial cell, MMP-9, claudin-5, PECAM-1, barrier integrity, ischemic injury

## Abstract

Endothelial integrity is critical in mitigating a vicious cascade of secondary injuries following acute ischemic stroke (AIS). Matrix metalloproteinase-9 (MMP-9), a contributor to endothelial integrity loss, is elevated during stroke and is associated with worsened stroke outcome. We investigated the FDA approved selective sphingosine-1-phosphate receptor 1 (S1PR1) ligand, ozanimod, on the regulation/activity of MMP-9 as well as endothelial barrier components (PECAM-1, claudin-5, and ZO-1) in human brain microvascular endothelial cells (HBMECs) following hypoxia plus glucose deprivation (HGD). We previously reported that S1PR1 activation improves HBMEC integrity; however, specific mechanisms underlying S1PR1 involvement in barrier integrity have not been clearly elucidated. We hypothesized that ozanimod would attenuate an HGD-induced increase in MMP-9 activity which would concomitantly attenuate the loss of integral barrier components. Male HBMECs were treated with ozanimod (0.5nM) or vehicle and exposed to 3h normoxia (21% O_2_) or HGD (1% O_2_). Immunoblotting, zymography, qRT-PCR, and immunocytochemical labeling techniques assessed processes related to MMP-9 and barrier markers. We observed that HGD acutely increased MMP-9 activity and reduced claudin-5 and PECAM-1 levels, and ozanimod attenuated these responses. In situ analysis via PROSPER, suggested that attenuation of MMP-9 activity may be a primary factor in maintaining these integral barrier proteins. We also observed that HGD increased intracellular mechanisms associated with augmented MMP-9 activation, however ozanimod had no effect on these targeted factors. Thus, we conclude that ozanimod has the potential to attenuate HGD mediated decreases in HBMEC integrity in part by decreasing MMP-9 activity as well as preserving barrier properties.

**Graphical Abstract:** 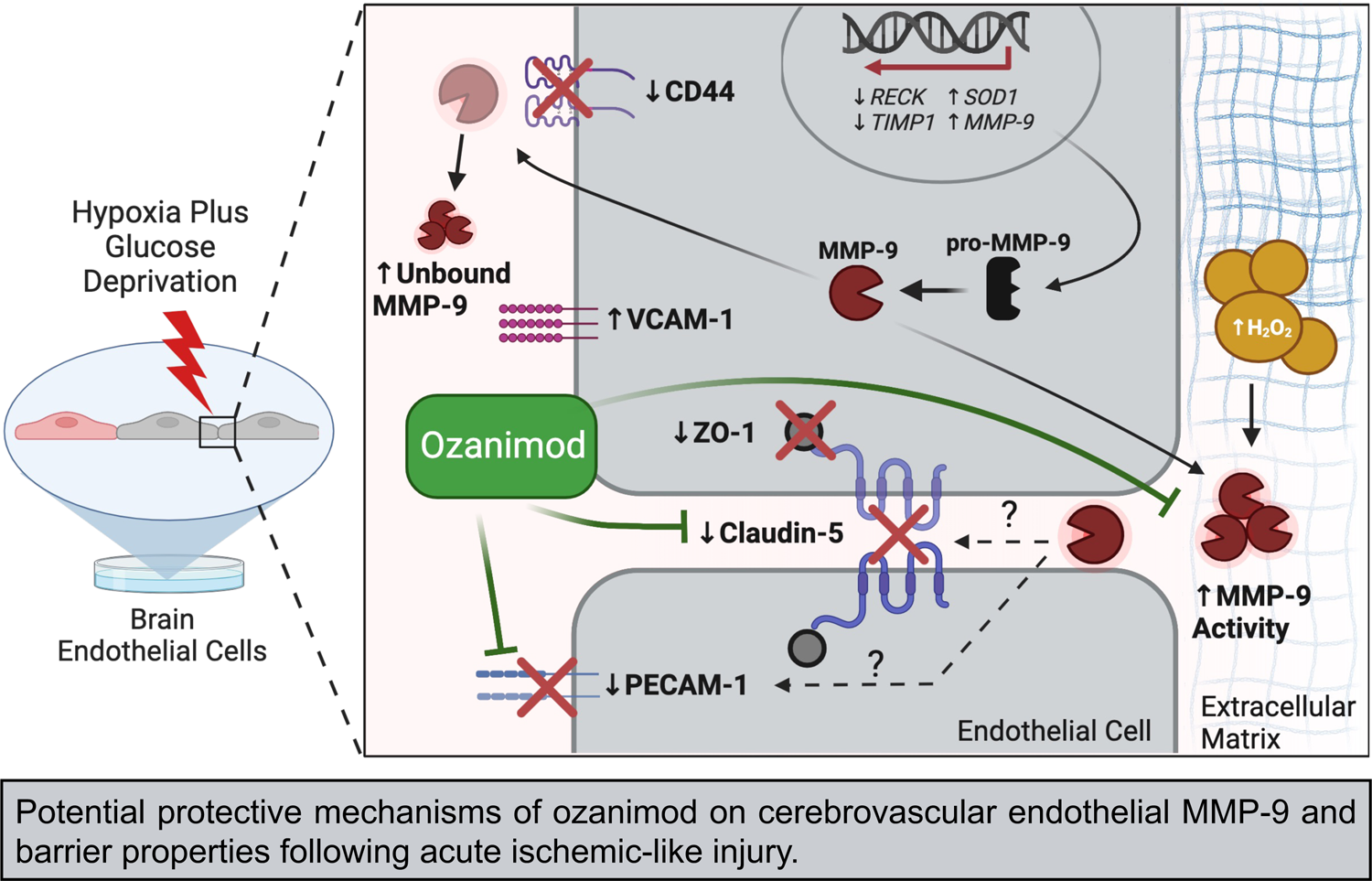

## INTRODUCTION

Matrix metalloprotease-9 (MMP-9) is a key enzyme involved in the complex and multimodal pathological vascular response during acute ischemic stroke (AIS). Upon endothelial activation, increased MMP-9 activity in-part, results in vascular integrity disruption and together can lead to a cascade of secondary injuries including barrier integrity loss, cerebrovascular inflammation, edema, and infarct expansion which all contribute to increased hemorrhagic risk and worsened stroke outcomes if the occlusion is not resolved. (1–4) Maintenance of neuronal survival is dependent upon the proper function of the blood brain barrier (BBB) which at the interface with the central nervous tissue is comprised of endothelial cells, astrocytes, pericytes, neurons, and extracellular matrix (ECM). [Reviewed in (5)]. Of these cells, the endothelium are primary regulators of permeability and if perturbed contributes significantly to the pathophysiological progression of AIS. (6, 7) At the level of the endothelium, a portion of its regulatory role is attributed to specific tight junction proteins and adhesion molecules such as claudin-5 (8), zonula occludens 1 (ZO-1) (9), platelet endothelial cell adhesion molecule 1 (PECAM-1) (10), and CD44 (11) which comprise critical aspects of barrier function and subsequent regulation of extracellular trafficking. In addition to tight junctions, the endothelium within the neurovascular unit has been shown to secrete ECM which plays an integral role in maintaining vessel and BBB integrity. [Reviewed in (12)] During the progression of AIS, tight junctions (13–17) and ECM are disrupted (18–22), which in part can be contributed to the notable acute activation of MMP-9. (23–26) Moreover, MMP-9 is postulated to be one of the drivers responsible for the biphasic manner in which the BBB is disrupted, where the acute onset phase is classified as detrimental, and the late onset is beneficial in promoting vascular remodeling. (26–29) Targeting of MMP-9 activity during the acute phase of BBB opening may serve as a likely potential therapeutic target to improve functional outcomes following ischemic injury. (30)

In efforts to mitigate worsened stroke outcomes, the development of synthetic compounds targeting selective sphingosine-1-phosphate receptors (S1PRs) as a novel approach for stroke treatment has gained significant traction. (31–36) Ozanimod, an FDA approved compound, is used to manage multiple sclerosis (37) and ulcerative colitis. (38) This selective S1PR1 ligand has been further explored as a potential therapeutic agent for stroke patients and has shown beneficial effects on improving cerebrovascular integrity, neurological function, and reducing infarct volume following ischemic injury. (39–43) S1PRs are a family of five G-protein coupled receptors (S1PR1-5) connected with an array of intracellular second messengers and signaling pathways. S1PR1 is a G_i_ protein (44) that ubiquitously and differentially expressed across various tissues, and has been linked to multiple pathways including PI3K/Akt, (45) Rac activation, (46) and MAPK. (47) Although studies have shown that S1PR ligands contain lymphocyte modulatory properties, S1PR1 agents can directly elicit effects at the level of the vasculature. This is supported by the findings that S1PR1 protein is expressed in vascular smooth muscle (48) as well as endothelial cells. (49) In addition to expression, a study by Grailhe et al. demonstrated maintenance of electrical impedance via ozanimod, in a human umbilical vein endothelial cell line. (50) Moreover, it has been observed that activation of endothelial S1PR1 promotes vascular integrity, (51–54) blood pressure homeostasis, (55) cell survival, (47, 56) and maintenance of microvascular patency and perfusion following ischemic stroke. (41) These functional outcomes secondary to endothelial S1PR1 activation showcase the potential of targeting S1PR1 to attenuate infarct expansion, reduce occurrence of hemorrhagic transformation, and mitigate stroke injury. (41)

To the best of our knowledge the role of S1PR1 on MMP-9 activity using a stroke model has not been investigated. We have however previously demonstrated that ozanimod attenuates increased brain endothelial cell barrier permeability via S1PR1 activation following acute exposure to hypoxia plus glucose deprivation (HGD), an in vitro model of ischemic injury; (40) however, specific mechanisms underlying S1PR1 involvement on barrier integrity have not been clearly elucidated. Therefore, in this study, we investigated a potential underlying mechanism and functional outcomes behind the HGD-induced endothelial cell permeability. Additionally, we addressed the beneficial potential of ozanimod on attenuating HGD-mediated alterations in paracellular permeability related structures such as tight junctions, adhesion molecules, and MMP-9 activity as well as underlying MMP-9 activation processes. We hypothesized that ozanimod would attenuate an HGD-induced increase in MMP-9 activity which would concomitantly attenuate the loss of integral human brain endothelial barrier components related to endothelial barrier permeability.

## METHODS

### Human Brain Microvascular Endothelial Cell Culture

Vendor-purchased cryopreserved subcultures of primary male human brain microvascular endothelial cells (HBMECs; Cell Systems Kirkland, WA, Catalogue number: ACBRI 376 Lot number: 376.02.03.01.2F) were received at *passage 3* and seeded at a density of 2 to 3 × 10^5^/cm^2^. HBMECs were cultured as previously described. (57) In brief, they were grown in 5% CO_2_ and room air at 37°C in phenol red-free Complete Classic Medium (10% FBS; fetal bovine serum) supplemented with Bac-off Antibiotic (Cell Systems, Catalogue number: 4Z0-643) and CultureBoost (Cell Systems, Catalogue number: 4CB-500). Cells were cultured in sterile 100mm × 20mm petri dishes (Corning; Catalogue number: 430591) which were all coated with Attachment Factor (Cell Systems; Catalogue number: 4Z0-201). Once cells reached 75-80% confluency, selected plates were cryopreserved for future use, and the remaining cells were continued in culture and studied at *passage 7*. For ICC and other cell imaging experiments, 100mm petri dishes were split and seeded on sterilized round cover glasses (Fisherbrand Scientific; Catalogue number: 12-545-100) treated with Attachment Factor within sterile 6-well plates (Corning; Catalogue number: 3516) and grown to 75-80% confluency.

### Hypoxia Plus Glucose Deprivation (HGD) Exposure and Drug Treatment

Ozanimod (RPC1063, 5-[3-[(1S)-2,3-dihydro-1-[(2-hydroxyethyl)amino]-1H-inden-4-yl]-1,2,4-oxadiazol-5-yl]-2-(1-methylethoxy)-benzonitrile), a selective sphingosine-1-phosphate receptor 1 (S1PR1) ligand (Cayman Chemical; Catalogue number: 19922), was prepared fresh under sterile conditions on the same day of administration for planned experiments as previously described. (48) Briefly, stock solutions (4mM) were made in dimethyl sulfoxide (DMSO) and further diluted in 1X Dulbecco’s phosphate buffer solution (DPBS) (Corning; Catalogue number: 21-031-CV) to a final working concentration (0.5nM). Similarly, W146 ([3R-amino-4-[(3-hexylphenyl)amino]-4-oxobutyl]-phosphonic acid, mono(trifluoroacetate)), a selective S1PR1 antagonist (Cayman Chemical; Catalogue number: 10009109), was freshly prepared under sterile conditions on the same day of experiments. Stock solutions (10mM) were made with grade methanol and further diluted in 1X DPBS to a final working concentration (10μM). This working solution and preincubation period was chosen as it has been reported that W146 at a concentration of 10μM elicits no off-target S1PRs (i.e., S1PR2, S1PR3, and S1PR5) agonist nor antagonist effects and has been previously utilized and shown to inhibit the effects of a highly selective S1PR1 ligand. (58) Vehicle solutions utilized in the control groups were generated in tandem and consisted of a DMSO/DPBS solution as well as a methanol/DMSO/DPBS solution prepared at a non-toxic concentration (DMSO <0.1%, methanol <2.0%). These solutions were maintained at 37°C during preparation and administration to HBMEC cultures. Prior to experimentation, designated plates were divided into the following groups: normoxia or HGD. To evaluate the effects of ozanimod, designated plates were divided into the following groups: normoxia plus vehicle (Normoxia + Vehicle), normoxia plus ozanimod (Normoxia + Ozd), HGD plus vehicle (HGD + Vehicle), or HGD plus ozanimod (HGD + Ozd). To test whether S1PR1 dependence, the following groups were added: normoxia plus ozanimod plus W146 (Normoxia + Ozd + W146) and HGD plus ozanimod plus W146 (HGD + Ozd + W146). Prior to HGD exposure, Complete Classic Medium was replaced with modified DMEM (ThermoFisher Scientific; Catalogue number: A14430-01) media containing no L-glutamine, phenol red, sodium pyruvate, or D-glucose. Next, cells were pretreated with ozanimod or vehicle, and in some select plates the antagonist, W146, was administered 0.5h prior to ozanimod treatment. Following drug treatment, plates designated for HGD were placed in a hypoxic sub chamber (BioSpherix) at 37°C and flushed with a medical grade gas mixture of 1% O_2_, 5% CO_2_, and nitrogen balance and exposed for 0.5-3h depending on the experimental design. Normoxic control plates containing Complete Classic Medium were replaced with fresh medium and incubated in a separate CO_2_ incubator at 37°C at 21% O_2_.

### Cellular Protein Isolation and Analysis

The isolation and quantification of protein performed is previously described. (48) Briefly, culture plates were removed from either the hypoxic BioSpherix chamber or normoxic incubator, placed on ice, washed twice with ice-cold phosphate buffered solution (PBS), trypsinized (0.25%) for 3min at 37°C (ThermoFisher Scientific; Catalogue number: 15090046) which was then neutralized with trypsin neutralizer (ThermoFisher Scientific; Catalogue number: R002100), and pelleted (centrifuge: Thermo Sorvall Legend RT+) at 1,900g (10 min, 4°C). The pellet was rinsed with ice-cold PBS, centrifuged, supernatant discarded, and pellet resuspended in ice-cold lysis buffer solution (50mM Tris-HCl, 150mM NaCl, 0.1% sodium dodecyl sulfate, 1% Nonidet P-40) containing protease inhibitors (20μM pepstatin, 20μM leupeptin, 0.1U/mL aprotinin, 0.1mM PMSF, 1mM DTT), phosphatase inhibitors (25mM β-glycerophosphate, 0.1mM sodium orthovanadate), 1mM EDTA, and 1mM EGTA for 15min. The resulting solution was then homogenized, sonicated, and centrifuged (Fisher Accuspin Micro 17R) at 17,000g (10 min, 4°C). Following centrifugation, the supernatant (whole cell lysate) was collected and stored at −80°C. Total protein was determined on the day of western blot experimentation and determined by using a bicinchoninic acid protein assay kit (ThermoFisher Scientific; Catalogue number: 23227) according to manufacturer’s protocol and measured on a Safire II (Tecan) plate reader.

### Western Blot

Protein levels for MMP-9 as well as for endothelial barrier and activation markers were examined following normoxic or HGD exposure using standard western blotting described previously. (48, 59) Briefly, sample lysates were diluted in Tris-glycine SDS sample buffer with the reducing agent, β-mercaptoethanol (10%) and boiled for 5min. The equally diluted samples and fluorescent standards (LI-COR Biosciences) were loaded onto 4–15% gradient Mini-PROTEAN TGX precast protein gels, (Bio-Rad Laboratories; Catalogue number: 4561084). Protein separation occurred by SDS-PAGE using a mini PROTEAN Tetra electrophoresis system (Bio-Rad Laboratories). The separated proteins were then transferred to nitrocellulose membranes using the same apparatus and membranes were then blocked in 3% nonfat dry milk rehydrated in Tris-Buffered saline containing 1% Tween (TBST; 20mM Tris, 150mM NaCl) at room temperature for 1h. Blots were then probed overnight at 4°C with MMP-9 (Cell Signaling; Catalogue number: 3852S), PECAM-1 (Sigma Aldrich; Catalogue number: SAB4502167), VCAM-1 (Santa Cruz Biotechnology; Catalogue number: sc-13160), claudin-5 (ThermoFisher Scientific; Catalogue number: 34-1600), or ZO-1 (ThermoFisher Scientific; Catalogue number: 33-9100) primary antibodies which were all diluted to 1:500 in TBST. After TBST wash cycle (5×, 5min), membranes were incubated in goat anti-rabbit IRDye 800CW dye (LI-COR) or goat anti-mouse IRDye 800CW dye (LI-COR) secondary antibodies at room temperature for 1h. Following another 5×, 5min TBST wash cycle, protein bands were visualized using an Odyssey Classic infrared imager (LI-COR). Next, membranes were exposed to overnight at 4°C with 1:15,000 anti-β-actin (Sigma Aldrich; Catalogue number: A5441) and for 1h at RT goat anti-mouse IRDye 680LT (LI-COR) which served as a loading control. Band densities were then analyzed using ImageStudio 3.0 software (LI-COR).

### Polymerase Chain Reaction (PCR)

Reverse transcription PCR (RT-PCR) was used to determine the presence of *MMP-9*, *RECK*, and *TIMP1* mRNA in HBMEC cultures. Briefly, RNA was extracted using the Qiagen kit (Thermo Fisher, Cat. No. 74034) according to the manufacturer’s instructions. HBMECs were collected and centrifuged (Thermo Sorvall Legend RT +) at 3,000 rpm for 10min at 4°C. The subsequent cell pellets then underwent a series of washes and centrifugations per the manufacturer’s instructions which resulted in collection of RNA in 14μL of diethyl pyrocarbonate (DEPC). Purity and concentration of extracted RNA was confirmed using a Nanodrop 2000 (ThermoFisher Scientific). RNA was reverse transcribed to make cDNA using the SuperScript III First-Strand Synthesis System (ThermoFisher Scientific; Catalogue number: 18080051) according to the manufacturer’s instructions. Each reaction mixture contained equal concentration loading of cDNA, as determined by cDNA concentration via Nanodrop 2000, 0.25μL of dNTP (ThermoFisher Scientific; Catalogue number: 18427088), 1μL 10× Taq Buffer (ThermoFisher Scientific; Catalogue number: EP0702), 0.25μL of Taq polymerase (ThermoFisher Scientific; Catalogue number: EP0702), 9.25μL of DEPC, 0.25μL of forward primer, and 0.25μL of reverse primer. All primers utilized in this study were designed in house utilizing the NCBI and UCSC genome libraries and obtained from Integrated DNA Technologies with standard desalting and at 25nM scale. The primer sequences used to detect *MMP-9* expression were 5’-GTCGTGGTTCCAACTCGGTTTG-3’ (forward) and 5’-GTGGTACTGCACCAGGGCAAG-3’ (reverse) and yielded an expected product size of 131bp. The primer sequences used to detect *RECK* expression were 5’-GCAGTGCGGGTGCATTGTGTTGTAATC-3’ (forward) and 5’-CAACCATTGTCTCTGGGCAATAATCTGGG-3’ (reverse) and yielded an expected product size of 147bp. The primer sequences used to detect *TIMP1* expression were 5’-CTTCTGCAATTCCGACCTCGTCATCAGG-3’ (forward) and 5’-CGAACCGGATGTCAGCGGCATC-3’ (reverse) and yielded an expected product size of 149bp. The reaction involved an initial melting step at 95°C for 3min followed by 35 cycles of 95°C (denature) for 30s, primer-specific annealing temperature for 30s, and 72°C (elongation) for 60s, then a 5min incubation at 72°C to terminate the reaction. Primer annealing temperatures for *MMP-9*, *RECK*, and *TIMP1* are 64.1°C, 65.6°C, and 67.4°C, respectively. Following the PCR reaction, samples were loaded alongside a UV standard (Bioland Scientific; Catalogue number: DM01-01) onto a 1% agarose gel. Gels were then run in a 1× tris-acetate-EDTA (TAE) (ThermoFisher Scientific; Catalogue number: B49) solution for 15min at 75V following by 35min at 95V. Gels were finally visualized and imaged using a ChemiDoc XRS UV gel imager to confirm the presence or absence of the mRNA of interest.

### Quantitative real time PCR

Quantitative real time PCR (qRT-PCR) was utilized to measure changes in mRNA levels of *MMP-9*, *RECK*, *TIMP1*, *SOD1*, and *SOD2*. RNA extraction and generation of cDNA was performed as described in the previous *polymerase chain reaction* section. All primer efficiencies were assessed via serial dilution from 200ng/μL to 0.02ng/μL and were found to be within 90-110% efficient. Following cDNA quantification via Nanodrop 2000 (ThermoFisher Scientific), cDNA from each sample was diluted to a working solution of 50ng/μL in DEPC. Duplicate wells were used for each target within the 96-well plate (USA Scientific; Catalogue number: 1402-9100). Within each well a mixture of cDNA (5μL), target primers diluted in DEPC (5μL; 0.4μM forward primer, 0.4μM reverse primer), and POWER SYBR Green PCR Master Mix (ThermoFisher Scientific; Catalogue number: 4368706) was added in order. The 96-well plates were then centrifuged (Thermo Sorvall Legend RT+) at 1,000g for 2min and then immediately loaded into the QuantStudio 6 and 7 Flex real-time PCR system (ThermoFisher Scientific). A two-step cycling protocol was implemented to collect cycle threshold (Ct) values of specific target amplicons and housekeeping mRNA *GAPDH* using the ΔΔCt relative quantification setting within QuantStudio 6. The reaction involved 2min at 50°C, an initial denaturing step at 95°C for 10min followed by 40 cycles of 95°C (denature) for 15s and primer-specific annealing temperature for 1min. Primer annealing temperatures for *MMP-9*, *RECK*, *TIMP1 SOD1*, *SOD2*, and *GAPDH* are 64.1°C, 65.6°C, 67.4°C, 62.4°C, 64.6°C, and 64.4°C respectively. The primer sequences used to detect *MMP-9*, *RECK*, and *TIMP1* expression are described in the previous *polymerase chain reaction* section. The primer sequences used to detect *SOD1* were 5’-GGAGATAATACAGCAGGCTGTACCAG-3’ (forward) and 5’-CCACACCATCTTTGTCAGCAGTCAC-3’ (reverse). The primer sequences used to detect *SOD2* expression were 5’-GTTTTGGGGTATCTGGGCTCCAG-3’ (forward) and 5’-GGTGACGTTCAGGTTGTTCACGTAG-3’ (reverse). The primer sequences used to detect *GAPDH* were 5’-GGTGTGAACCATGAGAAGTATGACAACAGC-3’ (forward) and 5’-CCTTCCACGATACCAAAGTTGTCATGG-3’ (reverse).

### Immunocytochemistry

Immunocytochemical assessment of target proteins was previously described by our lab(48) and implemented in this study to assess CD44. Briefly, HBMECs at density of 3 to 4 × 10^5^/cm^2^ were seeded onto glass coverslips coated with attachment factor (Cell Systems; Catalogue number: 4Z0-201) within 6-well plates at *passage 7*, grown to 70-80% confluency, and divided into groups as described before: normoxia plus vehicle (Normoxia + Vehicle), normoxia plus ozanimod (Normoxia + Ozd), HGD plus vehicle (HGD + Vehicle), or HGD plus ozanimod (HGD + Ozd). To test whether observed effects of ozanimod were S1PR1 dependent, some experiments included the following groups: normoxia plus ozanimod plus W146 (Normoxia + Ozd + W146) and HGD plus ozanimod plus W146 (HGD + Ozd + W146). Following 3h exposure/treatment, plates were immediately removed from incubator/sub chamber, placed on ice, media aspirated, and cells washed once with ice-cold DPBS. HBMECs were then fixed with 4% formaldehyde (10 min, RT) and subsequently washed twice for 4min with DPBS. HBMECs were then permeabilized with 0.1% TritonX-100 (Catalogue number: T9284) for 10 min and then blocked in 2% BSA (15min, RT). CD44 expression and nuclear localization was determined using anti-CD44 (Cell Signaling; Catalogue number: 3570S) and PECAM-1 expression using anti-PECAM-1 (Sigma Aldrich; Catalogue number: SAB4502167) at concentrations 1:100 in 2% BSA. Following an overnight incubation at 4°C, HBMECs were washed by 2% BSA (3×, 5min) followed by secondary antibody incubation with either Alexa Fluor 488 goat anti-mouse (ThermoFisher Scientific; Catalogue number: A11001) or Alexa Fluor 555 goat anti-rabbit (Thermo Fisher Scientific; Cat. No. A21428) at 1:6,000 in 2% BSA, for 1h at RT protected from light. A quick wash was then performed with the 2% BSA following by washes with DPBS (3×, 5min). Two coverslips from each treatment group were incubated in secondary antibody only and served as negative controls. Next, excess DPBS was removed, and the coverslips were mounted onto slides utilizing vectashield mounting medium with DAPI (Vector Laboratories; Catalogue number: H-1200). Coverslips were then sealed onto slides using nail polish and images captured with an inverted digital microscope (Keyence; Catalogue number: BZ-X800) following by area and localization quantification using ImageJ software.

### ImageJ Analysis Immunocytochemical Images

FIJI (NIH), an enhanced version of ImageJ2, was utilized to perform analysis on the area of CD44 and PECAM-1 normalized to the number of HBMECs as well as nuclear CD44 overlap using images from immunocytochemical experiments. Assessment of immunocytochemical images via FIJI, again has been previously described and implemented in this study with slight modifications. (48) Briefly, original immunocytochemical images obtained from the automated inverted digital microscope were inputted into FIJI and duplicated. The duplicate images were then split from RGB into three individual 8-bit images corresponding to the original red, green, and blue channels. From these 8-bit images, binary images were generated with a threshold individualized for either CD44, PECAM-1 or DAPI. CD44 and PECAM-1 analyzation of each treatment group was subjected to the automated IJ_IsoData threshold setting whereas DAPI analyzation of each treatment group was subjected to the automated Huang threshold setting. Next, the measurement parameters were set and redirected to the corresponding channel of the original image and overlaid with the original image inputted with constraints to not include any objects on the edge of the image. The subsequent objects generated corresponded to either the labeled CD44, PECAM-1, or nuclei and from which the total area of CD44 and PECAM-1 signal was quantized as μm^2^ and normalized to the number of DAPI stained nuclei present. Additionally, the CD44 constrained binary images were utilized by the JACoP macro-plugin without Costes’ automatic threshold to obtain M1 and M2 coefficients which subsequently quantified the colocalization of CD44 overlap with the nucleus and the nuclear overlap with CD44 respectively.

### Zymography

Media was extracted from cultures following exposure/treatment: ±HGD and ±ozanimod. The media was briefly centrifuged at 1,000g for 5min to remove any cellular debris and the supernatant was removed utilized as conditioned media. Total protein within conditioned media was determined on the day of zymography experimentation and determined by using a bicinchoninic acid protein assay kit (ThermoFisher Scientific) according to manufacturer’s protocol and measured on a Safire II (Tecan) plate reader. Conditioned media samples were diluted in Tris-glycine SDS sample buffer and the equally diluted samples and fluorescent standards (LI-COR Biosciences) were loaded onto Novex 10% zymogram plus gelatin protein gels (ThermoFisher Scientific; Catalogue number: ZY00100BOX). Protein separation occurred by SDS-PAGE using a mini– PROTEAN Tetra electrophoresis system (Bio-Rad Laboratories) at 100V for 2h and held at 4°C. After electrophoresis, the gel was incubated in 1× zymogram renaturing buffer (ThermoFisher Scientific; Catalogue number: LC2670) for 0.5h at RT with gentle agitation. Zymogram renaturing buffer was decanted and replaced with 1× zymogram developing buffer (ThermoFisher Scientific; Catalogue number: LC2671) and allowed to equilibrate the gel for 0.5h at RT with gentle agitation. Following equilibration, developing buffer was decanted and gels washed with nano-pure H_2_O and replaced with fresh 1× zymogram developing buffer and developed for 24h at 37°C. Following developing, gels were briefly washed with nano-pure H_2_O and stained with Coomassie blue staining solution (2.5g Coomassie Brilliant blue R-250, 450ml 100% methanol, 100ml 100% acetic acid, 400ml nano-pure H_2_O) for 2h at RT with gentle agitation. Gels were then washed with nano-pure H_2_O and de-stained with Coomassie blue de-staining solution (450ml 100% methanol, 100ml 100% acetic acid, 400ml nano-pure H_2_O) for 40min at RT with gentle agitation. The area of gelatin degradation at approximately 82kDa and 63kDa was indicative of MMP-9 and MMP-2 activity respectively and appeared as distinct white bands. The intensities of both MMP-9 and MMP-2 were obtained by an Odyssey Classic infrared imager (LI-COR) and inverse bands densities were then analyzed using ImageStudio 3.0 software (LI-COR).

### Extracellular H_2_O_2_ Quantification

Conditioned media was obtained as previously described in *zymography* section and utilized to ascertain concentration of hydrogen peroxide (H_2_O_2_). The aqueous pierce quantitative peroxide assay kit (ThermoFisher Scientific; Catalogue number: 23280) was utilized per manufacturer’s instructions to measure levels of H_2_O_2_ across treatment groups as previously described. In brief, equal loading of conditioned media was added to 96-well plate followed by 200μL of reaction mixture (1/100 v/v of Reagent A to Reagent B) for 20min at RT with agitation. Absorbance was then measured on a Safire II (Tecan) plate reader at 595nm and colorimetric quantification of H_2_O_2_ was compared to standards. Each sample was run in triplicate.

### Reagents

All reagents were purchased from Sigma Aldrich (St. Louis, MO) unless otherwise noted.

### Data and Statistical Analysis

Western blot and zymography analysis consisted of running lysates from each treatment group, including vehicles, on the same western blot or zymogram gel for direct comparison and homogeneity of data. Band densities during western blot analysis were normalized to loading control β-actin band density and data expressed as fold change relative to control group (i.e., normoxia or normoxia + vehicle). Experiments were repeated for statistical analysis and data graphed using Prism 8 (GraphPad Software). Confirmation of normal distribution was achieved with a Shapiro-Wilk test. *F*-test was performed to confirm if data sets achieved equal variance among groups. For analysis of datasets that achieved a normal distribution, direct comparisons between two groups were made using an unpaired *t* test alone if equal variance was achieved or with Welch’s correction if equal variance was not achieved. Comparisons between three or more groups were made using either one-way or two-way ANOVA with Tukey’s multiple comparisons post hoc test. Datasets that did not achieve a normal distribution underwent direct comparisons via Mann-Whitney test. *P* < 0.05 was considered significant. Values are expressed as means ± SEM.

## RESULTS

### HGD increased MMP-9 mRNA expression was not altered by ozanimod

We assessed the mRNA expression of *MMP-9* in HBMECs following acute ischemic-like exposure and determined if ozanimod would alter mRNA expression during normoxia and HGD. **Figure 1A** illustrates a representative image of a 2% agarose gel demonstrating confirmation of *MMP-9* mRNA expression within the HBMEC culture under 3h of normoxic control conditions or HGD. We observed that under normoxic conditions there was no detectable *MMP-9* mRNA expression, however HBMECs exposed to HGD revealed detectable amplicons. These results were then substantiated following qRT-PCR analysis of the *MMP-9* mRNA graphed in **Figure 1B**. Although, there was no detectable *MMP-9* mRNA expression with a 40 replication PCR cycle under normoxic conditions, there was a significant increase in *MMP-9* mRNA following HGD exposure. It was noted that HBMECs treated with ozanimod during HGD elicited greater variability in *MMP-9* mRNA expression compared to HGD alone. There was no difference between the two groups, suggesting that ozanimod does not alter acute HGD-mediated increases in *MMP-9* at the transcriptional level at this time point.

**Figure 1.**
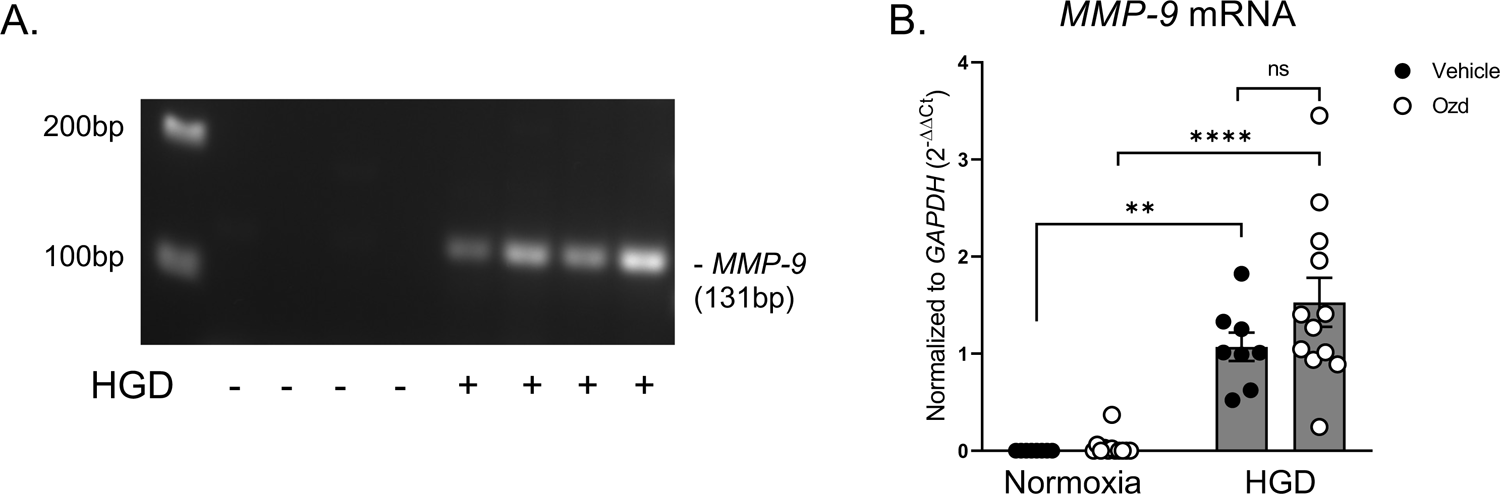
*MMP-9* mRNA levels in HBMECs following HGD in the presence or absence of ozanimod. (A) Representative RT-PCR gel for matrix metalloproteinase 9 (*MMP-9*) mRNA expression to verify expression in HBMECs under normoxia or HGD conditions that revealed presence of *MMP-9* only following HGD exposure. (B) Bar graph depicts qRT-PCR quantification of *MMP-9* mRNA present within HBMECs exposed to either normoxia or HGD and treated with either vehicle or ozanimod (Ozd; 0.5nM) normalized to the housekeeping mRNA (*GAPDH*) and expressed as 2^−ΔΔCt^. DMSO was used as the vehicle for the delivery of ozanimod. *n* = 8-12 (data are mean ± SEM). Two-way ANOVA with Tukey’s multiple comparisons post hoc test. Normoxia + Vehicle vs. HGD + Vehicle \*\**P* = 0.0011, Normoxia + Ozd vs. HGD + Ozd \*\*\*\**P* < 0.0001.

### Ratio of MMP-9 to pro-MMP-9 was increased following HGD and this response was not altered by ozanimod

Transcription and subsequent translation of MMP-9 leads to formation of a non-enzymatically active pro-MMP-9 which has a molecular weight of 92kDa. Upon activation, the pro-domain is removed resulting in a structure with a molecular weight of approximately 82-84kDa. (60) In these next set of experiments we determined whether HGD plus or minus ozanimod alters the intracellular HBMEC MMP-9 protein levels. **Figure 2A** illustrates a representative western blot using anti-MMP-9 to detect pro- and active forms of MMP-9 from HBMEC lysate. Densiometric analysis of intracellular MMP-9 revealed that HGD exposure induced a decrease in pro-MMP-9 (92kDa) when compared to the normoxic control, and ozanimod had no effect on this response (**Figure 2B**). In contrast, the enzymatically active form of MMP-9 (84kDa) following HGD exposure was not different compared to the normoxic control; however, there was a discernable increase in the variability of the protein levels when compared to the pro-MMP-9 expression following HGD exposure (**Figure 2C**). The impact of HGD in the conversion of pro-MMP-9 to MMP-9 was assessed via the individual ratio expression of pro vs. active for each sample. These data analysis demonstrated that HGD increased the ratio of active to inactive MMP-9 protein levels compared to normoxic controls; however, ozanimod did not attenuate this response (**Figure 2D**). These data viewed through the lens of previous data shown in **Figure 1** suggests that 3h HGD exposure may be upregulating the synthesis and then cleavage of MMP-9 which then becomes increasingly exported out of the HBMECs into the extracellular space.

**Figure 2.**
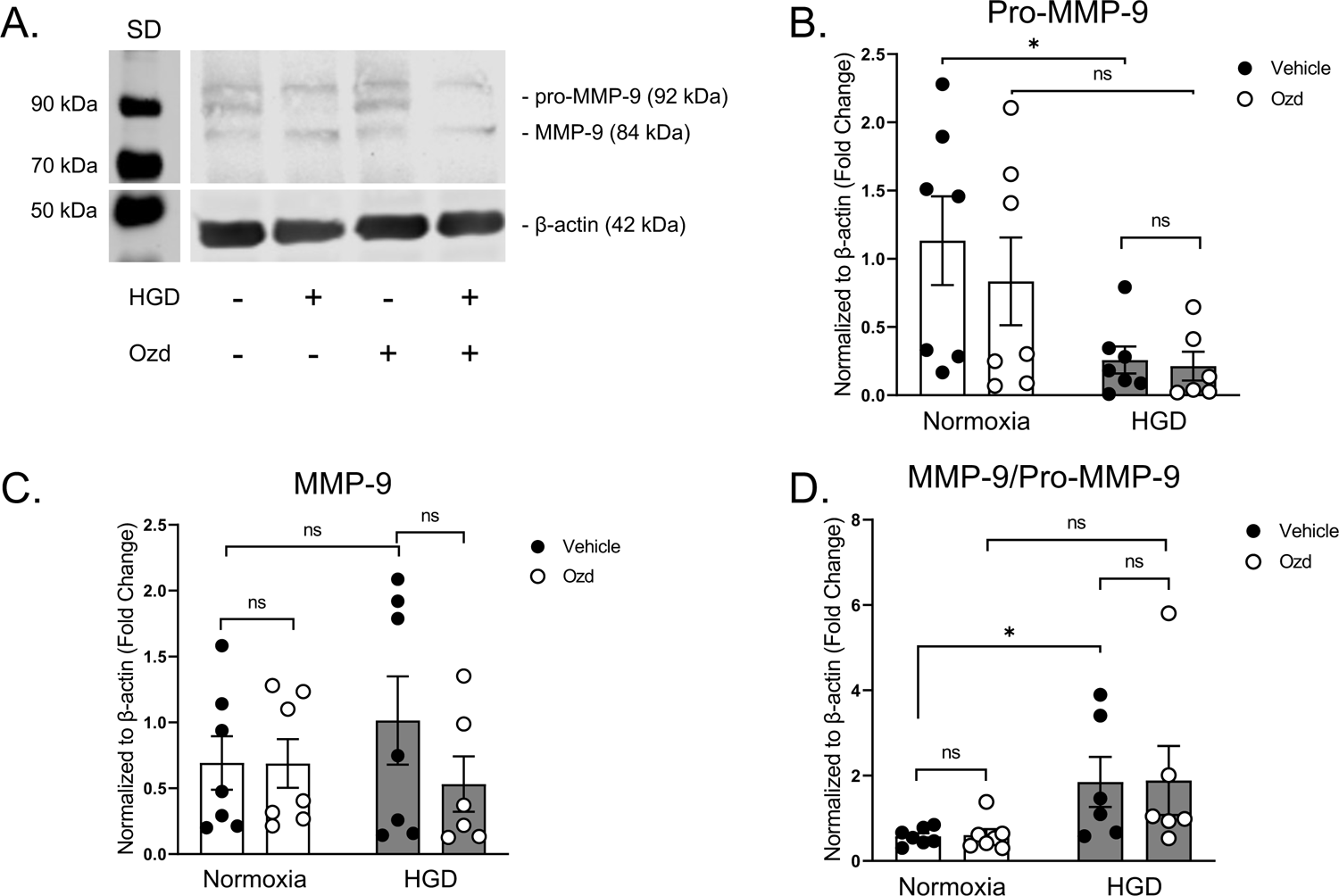
Intracellular HBMEC MMP-9 protein expression following HGD in the presence or absence of ozanimod. (A) Representative western blot illustrating the band migration for two distinct MMP-9 bands, pro-MMP-9 (92kDa; non-active form) and MMP-9 (84kDa; active form), as well as β-actin in HBMECs following treatment with either vehicle or ozanimod (Ozd; 0.5nM) and exposure to normoxia or HGD for 3h. Standard, SD. (B) Bar depicts pro-MMP-9 band intensities normalized to β-actin (loading control) and expressed as fold change ± SEM. *n* = 6-7 (data are mean ± SEM). Planned direct comparison was performed with unpaired *t* test with Welch’s correction. Normoxia + Vehicle vs. HGD + Vehicle \**P* = 0.0245. (C) Bar graph illustrates band intensities of MMP-9 normalized to β-actin (loading control) and expressed as fold change ± SEM. *n* = 6-7 (data are mean ± SEM). Two-way ANOVA with Tukey’s multiple comparisons post hoc test. (D) Bar graph demonstrates the ratio of MMP-9 (C) to pro-MMP-9 (B) intensities normalized to β-actin and expressed as fold change ± SEM. *n* = 6-7 (data are mean ± SEM). Planned direct comparison was performed with unpaired *t* test with Welch’s correction. Normoxia + Vehicle vs. HGD + Vehicle \**P* = 0.0392.

### HGD-induced increases in extracellular MMP-9 activity were attenuated by ozanimod

To address whether HGD alters HBMEC derived MMP-9 secretion and subsequent activity, we next assessed extracellular MMP-9 activity via zymography from conditioned media collected from HBMECs exposed to either normoxia or HGD in the presence or absence of ozanimod. A representative zymography image shown in **Figure 3A** demonstrates the band migration and enzymatic activity of both MMP-9 and MMP-2 from conditioned media of HBMECs. Inverse densiometric analysis of these bands are graphically represented in **Figure 3B** and **Figure 3C** respectively. MMP-9 activity was increased following HGD exposure and ozanimod reversed this response (**Figure 3B**). In contrast, there was no difference in MMP-2 activity following HGD when compared to the normoxic control (**Figure 3C**). These data demonstrate that acute HGD exposure increased HBMEC derived MMP-9 activity and that ozanimod treatment attenuated this response. Taken together, this suggests that although ozanimod had no effect at preventing synthesis of MMP-9 at the level of intracellular HBMEC mRNA and protein, the drug was effective at attenuating extracellular activity of the enzyme (**Figure 3D**). We next addressed how HGD exposure and treatment with ozanimod is eliciting these differential effects at an acute time point.

**Figure 3.**
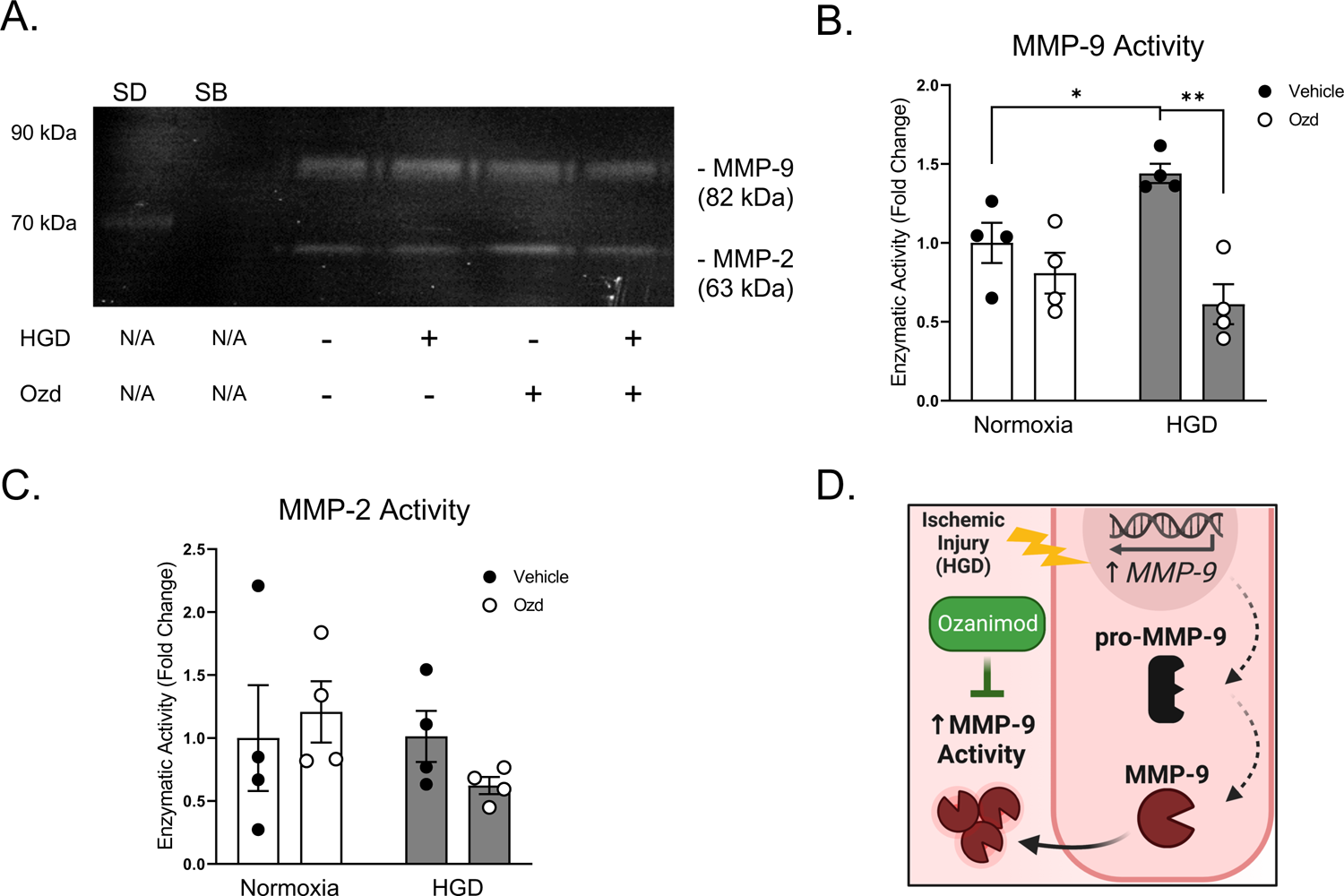
Extracellular HBMEC MMP-9 protein activity following HGD in the presence or absence of ozanimod. (A) Representative gelatin zymography illustrating the band migration and enzymatic activity for two distinct matrix metalloproteinase bands, MMP-9 (82kDa) and MMP-2 (63kDa), in conditioned media collected from HBMECs following treatment with either vehicle or ozanimod (Ozd; 0.5nM) and exposure to normoxia or HGD for 3h. Standard, SD; SB, Sample Buffer. (B) Bar depicts MMP-9 activity by inverse densiometric analysis and expressed as fold change ± SEM. *n* = 4 (data are mean ± SEM). Planned direct comparisons were performed with unpaired *t* test with Welch’s correction. Normoxia + Vehicle vs. HGD + Vehicle \**P* = 0.0323, HGD + Vehicle vs HGD + Ozd \*\**P* = 0.0013. (C) Bar graph illustrates quantification of MMP-2 activity by inverse densiometric analysis and expressed as fold change ± SEM. *n* = 4 (data are mean ± SEM). Two-way ANOVA with Tukey’s multiple comparisons post hoc test. (D) Graphical illustration of observed HBMEC MMP-9 regulation by HGD exposure and progression from intracellular mRNA and protein to extracellular activity and the observed attenuative effects of ozanimod on MMP-9 activity.

### HGD induced decreased in PECAM-1 levels was attenuated by ozanimod

PECAM-1, also known as CD31, has been shown to be cleaved by MMP-9 within the hepatic endothelium following ischemic injury. (16) Therefore, to validate the HGD-mediated increase in MMP-9 activity in HBMECs, PECAM-1 levels were assessed (**Figure 4**). Representative immunofluorescent anti-PECAM-1 labeling is illustrated in **Figure 4A**. Total area of anti-PECAM-1 signal normalized to the number of nuclei demonstrated that there was a decrease in labeled PECAM-1 protein following 3h HGD exposure and ozanimod attenuated this response (**Figure 4B**). When we examined PECAM-1 via immunoblotting we observed three distinct PECAM-1 bands (**Figure 4C**). The predicted molecular weight of PECAM-1 is approximately 80kDa which corresponds closest to the most prominent form of PECAM-1 detected in HBMECs, but the fully processed form corresponds to a molecular mass of 130kDa. (61) PECAM-1 has been described as forming homophilic and heterophilic interactions which are both implicated in the regulation of endothelial barrier integrity (62–64), thus suggesting that the distinct PECAM-1/PECAM-1 bands detected at 240-260kDa are PECAM-1 dimers while the detectable bands at 130kDa and 95kDa represent potential alternative splice forms of PECAM-1. (61) Further investigation into the regulation of potential alternative splicing or post-translational modification of PECAM-1 exceeds the scope of this study but opens the door for future studies to delineate mechanisms behind this observed alternative splicing. Taken together with our previous observation of ozanimod’s attenuative effect on MMP-9 activity (**Figure 3**), it is feasible to hypothesize that ozanimod may be sheltering PECAM-1 from MMP-9 mediated degradation and together suggests that ozanimod may be attenuating HGD-induced endothelial barrier disruption at the level of PECAM-1.

**Figure 4.**
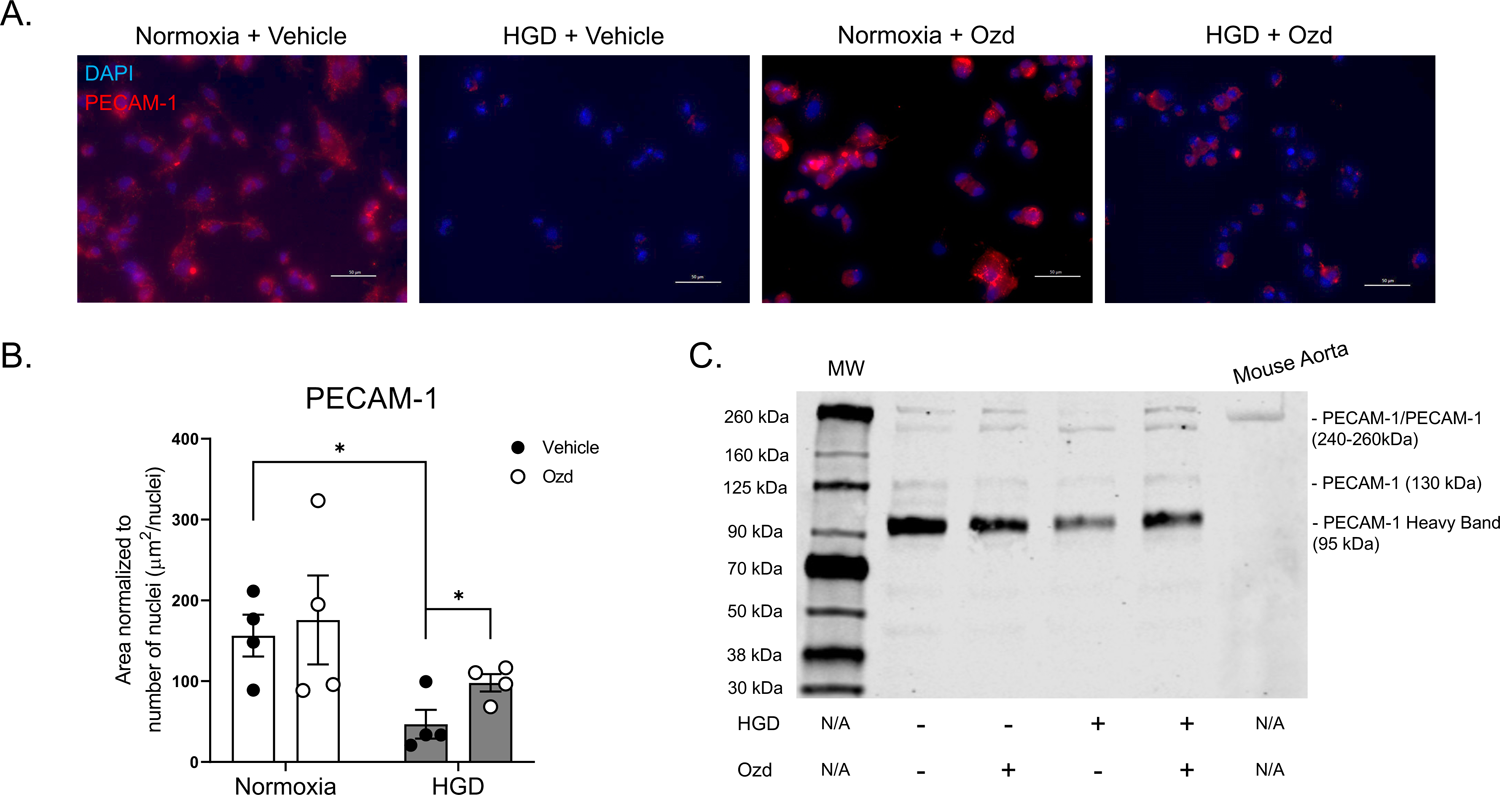
HBMEC PECAM-1 protein expression following HGD in the presence or absence of ozanimod. (A) Representative immunocytochemical images demonstrating the labeling of platelet endothelial cell adhesion molecule 1 (PECAM-1; red) in HBMECs following exposure to either HGD or normoxia for 3h and treatment with vehicle or ozanimod (Ozd; 0.5nM). Imaged at 40X magnification. Nucleus is labeled with 4’,6-diamidine-2’-phenylindole dihydrochloride (DAPI; blue). Scale bar, 50μM. (B) Bar graph depicts the relative quantification of PECAM-1 in HBMECs normalized to the number of nuclei and expressed as a ratio of the area occupied by PECAM-1 to number of nuclei (μm^2^/nuclei). *n* = 4 (data are mean ± SEM). Planned direct comparisons were performed with unpaired *t* test. Normoxia + Vehicle vs. HGD + Vehicle \**P* = 0.0131, HGD + Vehicle vs. HGD + Ozd \**P* = 0.0492. (C) Representative western blot illustrating the band migration for three distinct sets of platelet endothelial cell adhesion molecule 1 (PECAM-1) bands, PECAM-1/PECAM-1 (240-260kDa), PECAM-1 (130kDa), and a PECAM-1 Heavy Band (95kDa) in HBMECs following treatment with either vehicle or ozanimod (Ozd; 0.5nM) and exposure to normoxia or HGD for 3h. PECAM-1 expression was also demonstrated in mouse aortic tissue lysate to serve as a positive control for the PECAM-1 antibody but was not included in the interpretations of the study.

### Extracellular HBMEC derived H_2_O_2_ concentrations were increased following HGD exposure

To address how acute HGD exposure increased MMP-9 activity, we next examined levels of hydrogen peroxide (H_2_O_2_) which is exported extracellularly (65, 66) and has been shown to activate human endothelial MMP-9 after 2h. (67) We concomitantly choose to assess levels of vascular cell adhesion molecule 1 (VCAM-1) as it has been previously reported that VCAM-1, through an H_2_O_2_ mechanism, leads to endothelial MMP-9 activation. (68) We observed that HGD induced an increase in VCAM-1 levels in HBMECs compared to the normoxic controls and ozanimod did not alter this response (**Figure 5A**). We next examined mRNA levels of superoxide dismutase 1 (*SOD1*) and 2 (*SOD2*) and observed that HGD induced a significant increase in *SOD1* mRNA and ozanimod did not attenuate this response (**Figure 5B**). Our findings with *SOD1* mRNA levels contrasts with the assessment of *SOD2* mRNA levels, which revealed no difference across any treatment group (**Figure 5C**). We next examined the concentration of extracellular H_2_O_2_ within the conditioned media extracted from HBMEC cultures exposed to either normoxia or HGD and treated with either ozanimod or vehicle. There was an observed temporal dependent and HGD mediated increase in HBMEC derived extracellular H_2_O_2_ that peaked at 3h when compared to normoxic controls (**Figure 5D**). **Figure 5D** graphically illustrates extracellular H_2_O_2_ concentrations from conditioned media at time points 1 and 3h of HGD exposure. There was no difference in extracellular H_2_O_2_ at 1h; however, we did observe that HGD exposure significantly increased H_2_O_2_ at 3h both compared to the normoxic control as well as to the 1h HGD plus vehicle time point. These findings corroborate our previous findings of increased VCAM-1 protein and *SOD1* mRNA following 3h HGD exposure. Moreover, similar to our previous findings, we did not observe any effect of either ozanimod in either the normoxic or HGD exposure groups at any time point. There was also no discernible difference in extracellular H_2_O_2_ at 0.5h and 6h (data not shown) suggesting that H_2_O_2_ may be involved with an increase in MMP-9 activity. These data together suggest that the temporally dependent HGD mediated increases in VCAM-1 protein and *SOD1* mRNA may in part lead to an increase in extracellular H_2_O_2_ concentrations. Moreover, these elevated extracellular H_2_O_2_ concentrations may partially be responsible for the previously observed acute HGD mediated increase in MMP-9 activity however does not address the mechanism as to how ozanimod attenuated this response.

**Figure 5.**
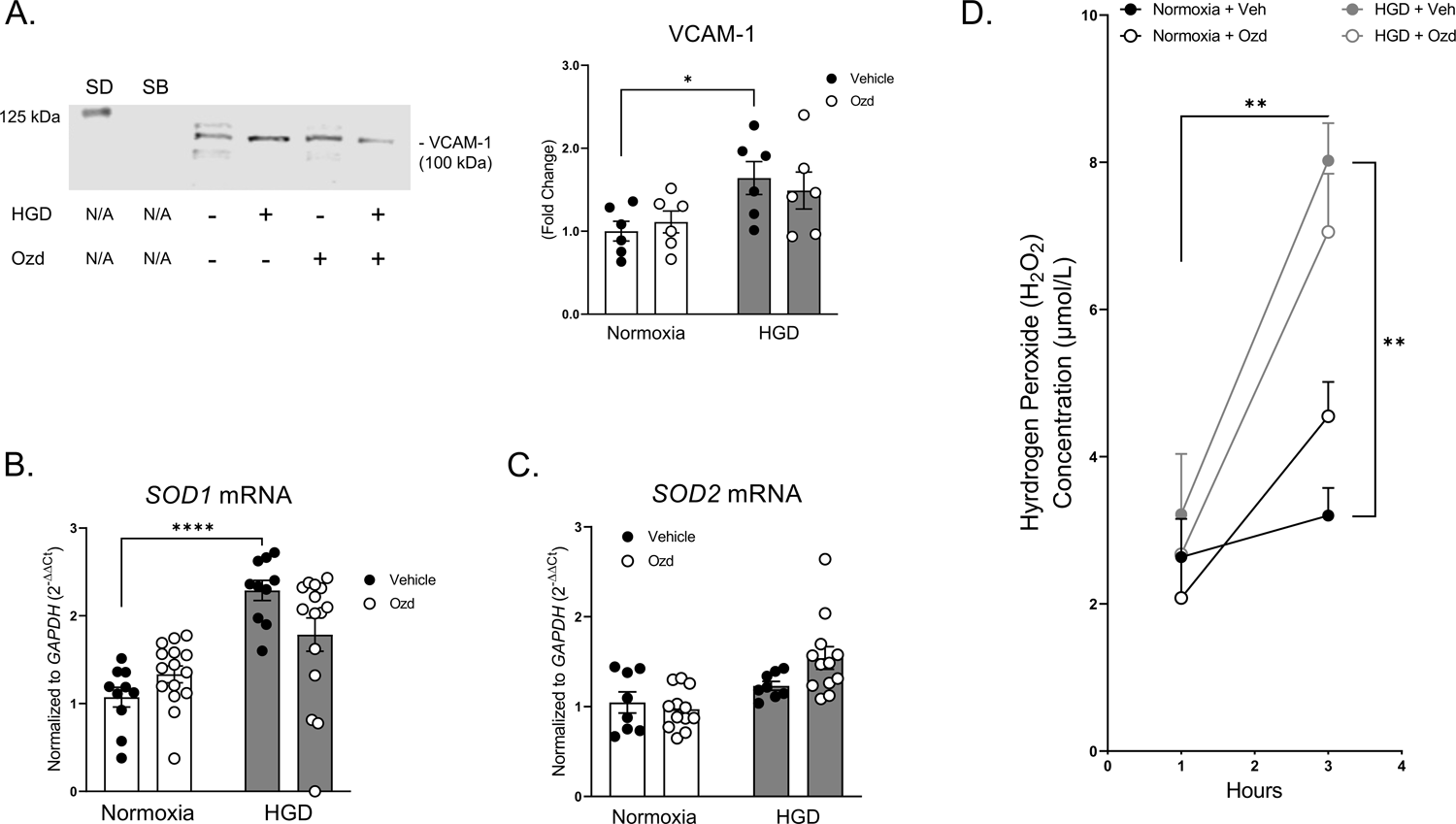
Extracellular hydrogen peroxide (H_2_O_2_) concentration derived from HBMECs following HGD in the presence or absence of ozanimod. (A) Representative western blot demonstrates the band migration of vascular cell adhesion molecule 1 (VCAM-1; 100kDa) in HBMECs following treatment with either vehicle or ozanimod (Ozd; 0.5nM) and exposure to normoxia or HGD for 3h. Bar graphs illustrate densiometric analysis of VCAM-1. *n* = 6 (data are mean ± SEM). Planned direct comparison was performed with unpaired *t* test with Welch’s correction. Normoxia + Vehicle vs. HGD + Vehicle \**P* = 0.0199. Standard, SD; SB, Sample Buffer. (B) Bar graph depicts qRT-PCR mediated quantitation of *SOD1* mRNA present within HBMECs exposed to either normoxia or HGD and treated with either vehicle or ozanimod (Ozd; 0.5nM) normalized to the housekeeping mRNA (*GAPDH*) and expressed as 2^−ΔΔCt^. *n* = 10-15 (data are mean ± SEM). Two-way ANOVA with Tukey’s multiple comparisons post hoc test. Normoxia + Vehicle vs. HGD + Vehicle \*\*\*\**P* < 0.0001, HGD + Vehicle vs. HGD + Ozd *P* = 0.0850. (C) Bar graph depicts qRT-PCR mediated quantitation of *SOD2* mRNA present within HBMECs exposed to either normoxia or HGD and treated with either vehicle or ozanimod (Ozd; 0.5nM) normalized to the housekeeping mRNA (*GAPDH*) and expressed as 2^−ΔΔCt^. *n* = 8-12 (data are mean ± SEM). Two-way ANOVA with Tukey’s multiple comparisons post hoc test. (D) Graph demonstrates the quantification of extracellular concentrations H_2_O_2_ from conditioned media extracted from HBMEC cultures exposed to either normoxia or HGD for 1 and 3h and treated with vehicle (Veh) or ozanimod (Ozd; 0.5nM). *n* = 3 (data are mean ± SEM). Two-way ANOVA with Tukey’s multiple comparisons post hoc test. 1h HGD + Vehicle vs. 3h HGD + Vehicle \*\**P* = 0.0015, 3h Normoxia + Vehicle vs. 3h HGD + Vehicle \*\**P* = 0.0014.

### HGD decreased inhibitory mechanisms of MMP-9 export and activation

Although we observed that HGD acutely increased mechanisms leading to MMP-9 activation, we did not observe an attenuative effect of ozanimod on these responses, therefore we next addressed potential mechanisms contributing to decreased MMP-9 activity. In human lung cancer cells, membrane anchored protein, reversion-inducing cysteine-rich protein with Kazal motifs (RECK), decreases MMP-9 secretion and activity that culminated in an attenuation of tumor migration (69, 70) suggesting that it could be acting in a similar manner during acute ischemic injury in the cerebrovasculature. Moreover, tissue inhibitors of metalloproteinases 1 (TIMP1) has been shown to inhibit almost all known MMPs with varying degrees of efficacy, but more specifically has been shown to associate with MMP-9. (71) Additionally, it has been previously observed that TIMP1 mRNA levels are in part dependent upon S1PR1. (72) Therefore, in this study we hypothesized that HGD exposure would decrease expression of RECK and TIMP1, which would be attenuated by treatment with ozanimod. **Figure 6A** demonstrates a representative PCR gel confirming *RECK* mRNA expression under both normoxic and HGD conditions, as well as that HGD decreased *RECK* mRNA expression. This was then confirmed using qRT-PCR measurement demonstrating a significant decrease in *RECK* mRNA following 3h HGD exposure; ozanimod did not alter *RECK* mRNA expression (**Figure 6A**). Similarly, *TIMP1* mRNA expression was visualized on an agarose gel that confirmed its expression profile in HBMECs following both normoxic and HGD exposure for 3h. Moreover, qRT-PCR analysis of *TIMP1* revealed a significant decrease in HBMECs exposed to HGD mRNA expression of the inhibitory molecule was not attenuated by ozanimod (**Figure 6B**). These data suggest, that in addition to increasing a potential mechanism of activation (**Figure 5**), acute HGD exposure decreased mechanisms related to inhibition of HBMEC MMP-9 activity; however, the previously observed functional effects of ozanimod on MMP-9 activity remains to be answered.

**Figure 6.**
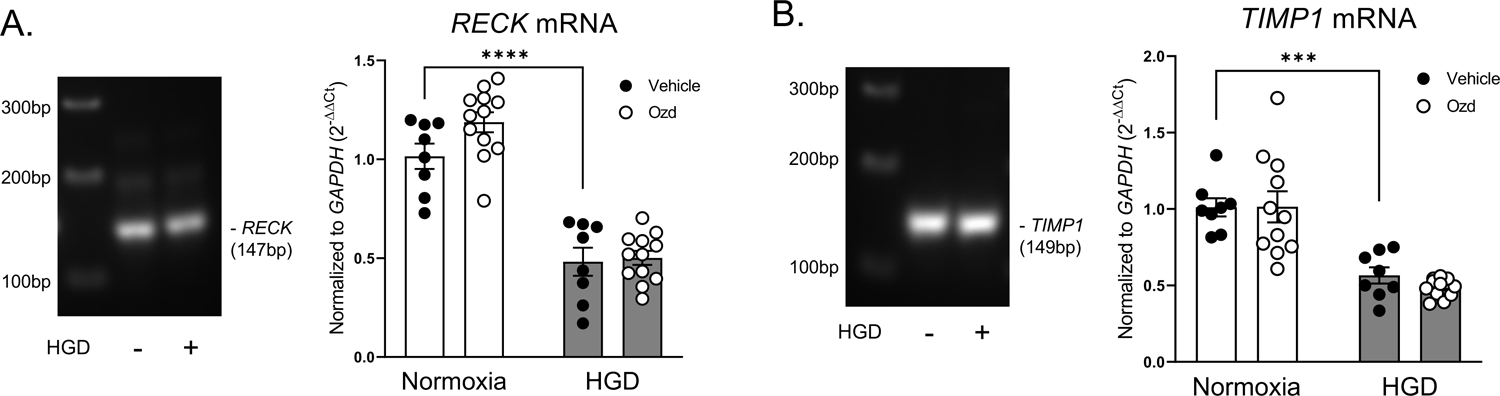
Extracellular transport, inhibition, and docking of MMP-9 in HBMECs following HGD in the presence or absence of ozanimod. (A) Bar graph depicts qRT-PCR mediated quantitation of *RECK* mRNA present within HBMECs exposed to either normoxia or HGD and treated with either vehicle or ozanimod (Ozd; 0.5nM) normalized to the housekeeping mRNA (*GAPDH*) and expressed as 2^−ΔΔCt^. *n* = 8-12 (data are mean ± SEM). Two-way ANOVA with Tukey’s multiple comparisons post hoc test. Normoxia + Vehicle vs. HGD + Vehicle \*\*\*\**P* < 0.0001. (B) Bar graph depicts qRT-PCR mediated quantitation of *TIMP1* mRNA present within HBMECs exposed to either normoxia or HGD and treated with either vehicle or ozanimod (Ozd; 0.5nM) normalized to the housekeeping mRNA (*GAPDH*) and expressed as 2^−ΔΔCt^. *n* = 8-11 (data are mean ± SEM). Two-way ANOVA with Tukey’s multiple comparisons post hoc test. Normoxia + Vehicle vs. HGD + Vehicle \*\*\**P* = 0.0008.

### HGD exposure decreased HBMEC CD44 levels

In efforts to identify a mechanism by which ozanimod might be acting to attenuate MMP-9 activity, we next utilized an immunocytochemical approach to ascertain the effects of HGD exposure as well as ozanimod on the levels and nuclear overlap of CD44 (**Figure 7**). Investigating the levels and distribution of CD44 was addressed to assess the functional component integral to MMP-9 activity as well as to endothelial barrier function. The multimodal cell surface protein, CD44, has been implicated in many cellular roles such as endothelial barrier function (11), proliferation and survival (73), and as an anchor point for MMP-9. (17, 74) **Figure 7A** illustrates representative immunocytochemical images of HBMECs labeled with anti-CD44 following normoxia or HGD exposure in the absence or presence of ozanimod. Total area of anti-CD44 signal was measured and normalized to the number of nuclei. There was a significant decrease in anti-CD44 following HGD when compared to the normoxic control and this response was not attenuated by ozanimod (**Figure 7B**). Intriguingly however, within the normoxic control groups we observed a noticeable impact of ozanimod on the distribution of CD44 with regards to the nucleus (**Figure 7C**). Quantification of the overlap of CD44 with the nucleus determined with Mander’s Coefficient via the JACoP macro-plugin within FIJI showed no difference between HBMECs exposed to normoxia and HGD, however treatment with ozanimod under normoxic conditions significantly reduced the overlap of CD44 with the nucleus. There was no difference in CD44 overlap with the nucleus in cells treated with ozanimod under HGD conditions. As part of Mander’s coefficient, we also obtained quantification of the inverse, which described the overlap of the nuclei with CD44 (**Figure 7D**). This analysis revealed no difference between HBMECs exposed to normoxia and HGD for 3h but demonstrated that ozanimod exhibited nearly identical effects with regards to overlap attenuation of the nuclei with CD44 under normoxic conditions (**Figure 7D**). The observed decreased levels of HBMEC CD44 following HGD exposure suggests a decrease in not only CD44 mediated barrier integrity, survival, and proliferation but also that secreted MMP-9 may be losing an anchor point from which to perform its enzymatic activity and thusly might be free to expand beyond the confines of the endothelial surface to perform degradation. Additionally, the observed decrease in CD44 overlap with the nucleus following ozanimod suggests reduced translocation of the intracellular domain of CD44 to the nucleus, which has been shown to induce transcriptional effects during cellular stress. (75–77) Since we observed that ozanimod did not attenuate HGD mediated decreases in CD44 but did previously reduce MMP-9 activity and preserved PECAM-1 expression, we hypothesized that untethered MMP-9 would negatively impact other contributors of endothelial barrier function and ozanimod would attenuate this response.

**Figure 7.**
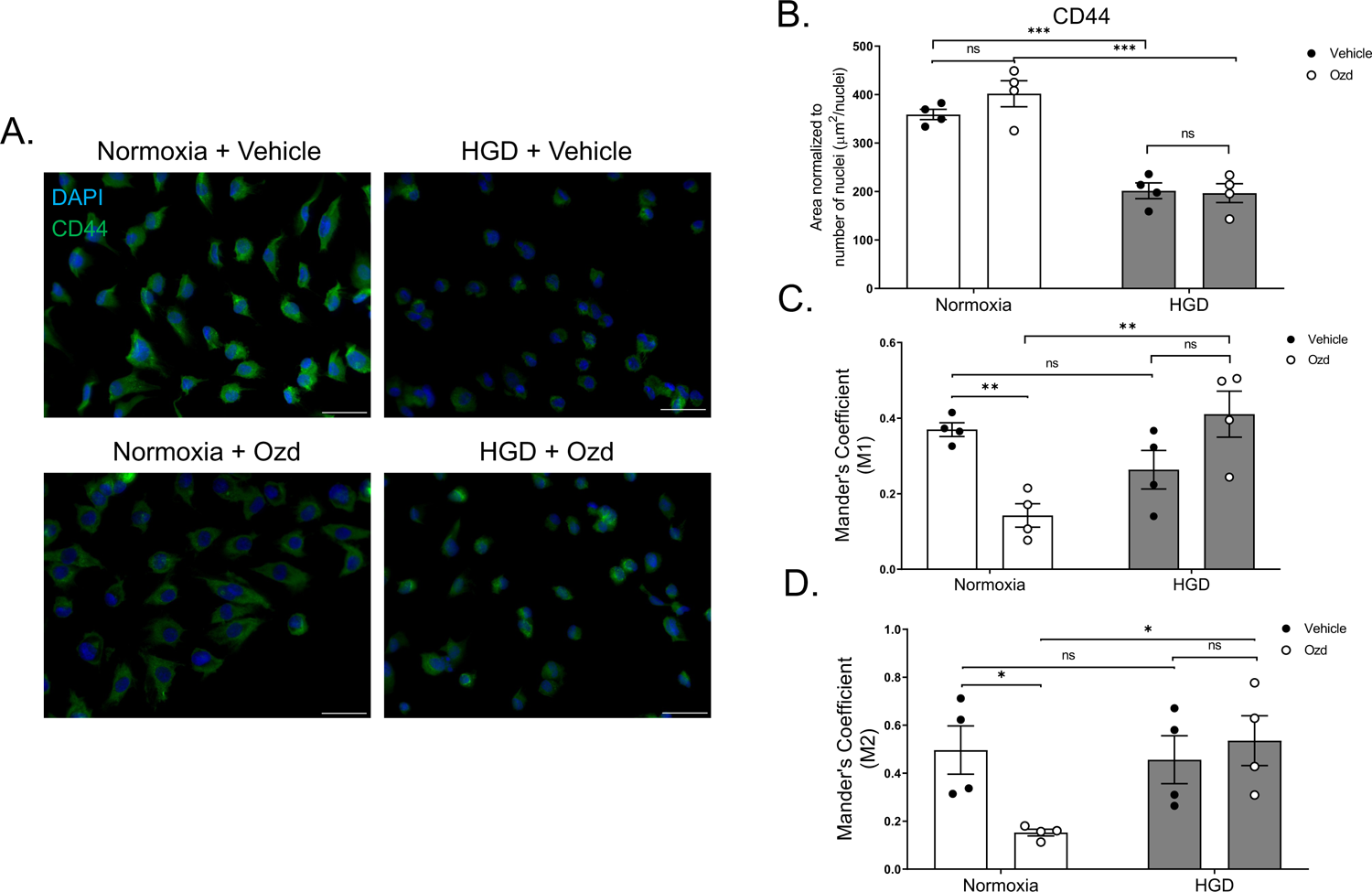
CD44 protein expression and nuclear localization in HBMECs following HGD in the presence or absence of ozanimod. (A) Representative immunocytochemical images demonstrating the nuclear or non-nuclear localization and labeling of anti-CD44 (CD44; green) in HBMECs following exposure to either HGD or normoxia for 3h and treatment with vehicle or ozanimod (Ozd; 0.5nM). Imaged at 40X magnification. Nucleus is labeled with 4’,6-diamidine-2’-phenylindole dihydrochloride (DAPI; blue). Scale bar, 50μM. (B) Bar graph depicts the relative quantification of CD44 in HBMECs normalized to the number of nuclei and expressed as a ratio of the area occupied by CD44 to number of nuclei (μm^2^/nuclei). *n* = 4 (data are mean ± SEM). Two-way ANOVA with Tukey’s multiple comparisons post hoc test. Normoxia + Vehicle vs. HGD + Vehicle \*\*\**P* = 0.0002. (C) Bar graph depicts Mander’s Coefficient (M1) of the overlap of CD44 with DAPI stained nuclei calculated from the images shown in (A). *n* = 4 (data are mean ± SEM). Two-way ANOVA with Tukey’s multiple comparisons post hoc test. Normoxia + Vehicle vs. Normoxia + Ozd \*\**P* = 0.0041, Normoxia + Ozd vs. HGD + Ozd \*\**P* = 0.0078. (D) Bar graph depicts Mander’s Coefficient (M2) of the overlap of DAPI stained nuclei with CD44 calculated from the images shown in (A). *n* = 4 (data are mean ± SEM). Two-way ANOVA with Tukey’s multiple comparisons post hoc test. Normoxia + Vehicle vs. Normoxia + Ozd \**P* = 0.0435, Normoxia + Ozd vs. HGD + Ozd \**P* = 0.0107.

### Ozanimod differentially attenuated HGD-mediated decreases in barrier proteins ZO-1 and claudin-5

Next, to test our hypothesis that the observed increased MMP-9 activity could functionally be involved in endothelial barrier dysfunction, we assessed protein levels of tight junction protein, claudin-5, and the intracellular anchoring protein integral for barrier function, zonula occludens 1 (ZO-1). We chose to assess these markers, as both claudin-5 and ZO-1 have been shown to be decreased acutely in murine brain endothelial cells following exposure to exogenous MMP-9. (13) **Figure 8A** demonstrates that HGD exposure induced a decrease in claudin-5 protein levels and ozanimod attenuated this response. Thus, suggesting that in addition to preserving PECAM-1 levels, ozanimod may be capable of preserving endothelial barrier integrity by reducing the loss of claudin-5 following an acute ischemic-like injury. Finally, immunoblotting of ZO-1 protein showed that HGD exposure significantly reduced the detectable levels of ZO-1 in HBMECs compared to the normoxic control; however, ozanimod treatment did not exert any effects on this response (**Figure 8C**). These data taken together with our previous observations that ozanimod attenuated an HGD-mediated reduction in trans-endothelial electrical resistance, (40) suggest that ozanimod, a selective S1PR1 ligand, may be functionally attenuating a loss in endothelial barrier integrity in part by attenuating extracellular MMP-9 activity and reducing the loss of PECAM-1 and claudin-5 proteins.

**Figure 8.**
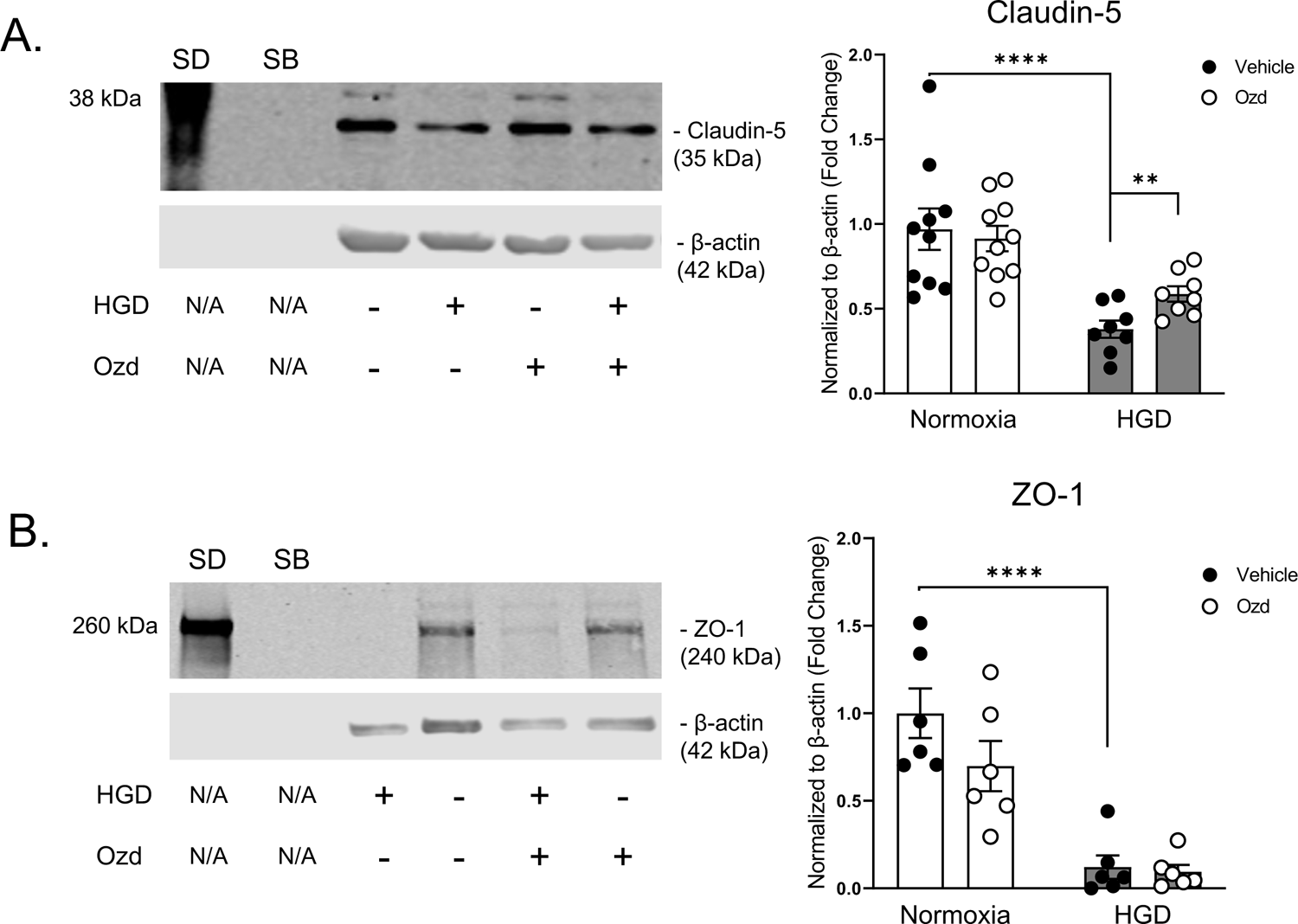
Levels of HBMEC barrier proteins ZO-1 and claudin-5 following HGD in the presence or absence of ozanimod. Representative western blots demonstrate the band migration of (A) claudin-5 (35kDa), (B) zonula occludens 1 (ZO-1; 240kDa), and β-actin (42kDa) in HBMECs following treatment with either vehicle or ozanimod (Ozd; 0.5nM) and exposure to normoxia or HGD for 3h. Bar graphs illustrate densiometric analysis of claudin-5 (A), and ZO-1 (B) expressed as fold change to normoxia plus vehicle. (A) *n* = 8-10 (data are mean ± SEM). Two-way ANOVA with Tukey’s multiple comparisons post hoc test. Normoxia + Vehicle vs. Normoxia + Ozd \*\*\**P* = 0.0002. Planned direct comparison was performed with unpaired *t* test with Welch’s correction. HGD + Vehicle vs. HGD + Ozd \*\**P* = 0.0093. (B) *n* = 6 (data are mean ± SEM). Two-way ANOVA with Tukey’s multiple comparisons post hoc test. Normoxia + Vehicle vs. HGD + Vehicle \*\*\*\**P* < 0.0001. Standard, SD; SB, Sample Buffer.

## DISCUSSION

In this study, using an in vitro ischemic-like stroke model, we have identified a potential novel mechanism by which ozanimod, a selective S1PR1 ligand, attenuates HGD-induced disruptions in human brain endothelial barrier function. To the best of our knowledge this is the first study to investigate the effect of a selective S1PR1 compound following ischemic-like conditions on MMP-9 regulation and endothelial barrier properties in a human brain endothelial cell model. To summarize, we demonstrate that HGD increased MMP-9 activation along with altered endothelial barrier proteins and that ozanimod may play a beneficial role in attenuating this HGD induced response. In brief, HGD exposure 1) induced MMP-9 transcription, 2) increased extracellular MMP-9 enzymatic activity, 3) enhanced levels of VCAM-1 and extracellular H_2_O_2_, 4) altered levels of tight junction, activation, and regulatory/anchoring proteins. We further demonstrated that ozanimod attenuated HGD induced MMP-9 activity as well as preserved protein levels of tight junction protein, claudin-5.

### MMP-9 mRNA and Protein Regulation

Maintenance of endothelial barrier integrity both during physiological conditions and following an ischemic event is important in the regulation of microvessel stability (78), blood flow (79), and ultimately blood brain barrier integrity. (80, 81) Enzymatic activity of MMP-9 is a key mediator of functional endothelial barrier disruption and serves clinically as a marker of ischemic stroke. (82, 83) Thus attenuation of MMP-9 activity remains an attractive therapeutic aim in the management of AIS, and here in this study we demonstrated that ozanimod attenuated extracellular MMP-9 activity following acute ischemic-like injury (**Figure 3B**). Complementary studies implementing genetic regulation of MMP-9 demonstrate reduced lesion volume and infarct expansion following focal ischemia. (84–86) These insights suggest that while genetic ablation of *MMP-9* may prove beneficial acutely, the long-term effect of such approaches may prove untenable. In addition, pharmacologic inhibition of MMP-9 with compounds such as BB-94 (87), KB-R7785 (88), melatonin (89), and fasudil (90) have been successful in pre-clinical animal stroke models; however, these therapeutics and others targeting MMPs have largely failed in translation to clinical application. This is exemplified by the utilization of minocycline which in pre-clinical animal models showed great promise (92), but was terminated following interim analysis during clinical trials due to determined futility (NCT00930020). Additional independent studies have suggested that while attenuation of MMP-9 during the acute phase of ischemic stroke is beneficial, this enzyme plays a key role in the later remodeling phase responsible for angiogenesis and subsequently reperfusion of the damaged area. (26, 93) This in part may be a reason for failure of previous pharmacological agents to transition from the pre-clinical model to clinical practice as they impacted both the detrimental acute and beneficial chronic phases of MMP-9 activity. A better understanding of regulation of MMP-9 levels and activity at the molecular level may provide better insight into therapeutic options for stroke. In vascular smooth muscle cells *MMP-9* is upregulated via NFκB activation (94), which could potentially be mediating the drastic increase in *MMP-9* mRNA following HGD exposure for 3h observed in this study (**Figure 1B**). Moreover, although levels of *MMP-9* mRNA were not changed with ozanimod during HGD exposure, we observed a noticeable increase in variability between samples. This could thusly be attributed to intercellular heterogeneity within the individual samples and between different cultures and potentially be due to differential regulation of NFκB. This postulation is supported by a study conducted in human pulmonary endothelial cells which showed that overexpression of S1PR1 decreased p65 phosphorylation and translocation to the nucleus. (95) Therefore, we conclude that 3h HGD exposure significantly upregulates *MMP-9* mRNA levels which sets the stage for a potential increase in protein production.

The process of MMP-9 translation is tightly regulated and has previously been shown to be regulated by phosphorylation of eukaryotic initiation factor 4E (96) which when pharmacologically attenuated resulted in reduced *MMP-9* translation. (97) In this along with HGD induced increases in levels of *MMP-9* mRNA (**Figure 1**), we noted a decrease in the levels of pro-MMP-9 (**Figure 2B**). From these findings we hypothesized that either the translation of *MMP-9* mRNA to pro-MMP-9 was attenuated or that the inactive pro-MMP-9 protein was being increasingly converted to the enzymatically active MMP-9 form. We observed an increase in the ratio of the active MMP-9 to inactive pro-MMP-9 protein levels (**Figure 2D**) following HGD exposure which further confounded the potential variables at play and suggests that HGD exposure either induced 1) a decrease in pro-MMP-9 synthesis or 2) a conversion rate of pro-MMP-9 to MMP-9 that balances the rate by which MMP-9 is being transported extracellularly to ultimately match the levels of endogenously expressed MMP-9 protein under normoxic conditions. In support of the second potential conclusion, it has been previously reported that human umbilical vein endothelial cells exposed to angiogenic factors experienced increased rapid vesicle formation and secretion that contain the proteolytically active enzyme, MMP-9, which appears to maximize in the extracellular space at 3h. (98) Additionally, we have observed a significant increase in HBMEC vesicle formations following exposure to 3h of HGD compared to normoxic controls (Wendt and Gonzales, unpublished observation 2023). To further address the questions posed by these findings of whether translation of *MMP-9* mRNA was being attenuated or if MMP-9 was being exported extracellularly following HGD exposure, as well as the role of ozanimod, we assessed extracellular MMP-9. Utilizing zymography, we observed increased extracellular MMP-9 activity following HGD exposure and that ozanimod attenuated this response (**Figure 3B**). Although we observed an increase in endothelial derived MMP-9 activity extracellularly, we detected no difference in MMP-2 activity (**Figure 3C**). These data together thusly suggest that 3h ischemia-like injury induces an increase in HBMEC MMP-9 transcription, protein exportation, and extracellular activity and treatment with ozanimod attenuates the functional extracellular enzymatic activity of MMP-9 (**Figure 3D**).

### MMP-9 Activation

In efforts to identify a potential mechanism by which 3h HGD exposure induced MMP-9 activity and by which ozanimod attenuated this response, we first measured VCAM-1 as it is a well-accepted marker of endothelial cell activation and has been implicated upstream of MMP-9 activation through production of H_2_O_2_. (68, 99, 100) Clinically, elevated intracranial levels of VCAM-1 correlated with increased infarct and edema volumes in stroke patients that underwent mechanical thrombectomies. (101) Moreover, Yousef et al demonstrated that both anti-VCAM-1 antibodies and genetic ablation of brain endothelial cell VCAM-1 reduced age-related neurodegeneration in part through reduced microglial reactivity and cognitive defects. (102) These studies highlight the significance of VCAM-1 in mediating both MMP-9 activation as well as mediating negative neurological outcomes. In our study we observed a significant increase in VCAM-1 protein levels following acute HGD exposure; however, there was no effect of ozanimod (**Figure 5A**). Previous work has demonstrated that translocation of active NFκB to the nucleus plays a critical role in the rapid induction of VCAM-1 in response to endothelial activation as well as inflammation. (103) This in part supports the previous hypothesis that HGD exposure induced an increase in NFκB activation. In addition to VCAM-1, we assessed mRNA levels of *SOD1* and *SOD2* following HGD exposure in the presence or absence of ozanimod. Both enzymes are well known and accepted for the ability to convert superoxide (O_2_^-^) to H_2_O_2_ [Reviewed in (104)] and in this study were hypothesized to contribute to the activation of MMP-9. Unexpectedly and counter to its consideration as a “housekeeping gene”, we observed a significant increase in mRNA levels of *SOD1* following HGD (**Figure 5B**). Although, *SOD1* shares significant overlap in transcription factors associated with regulation of both *SOD2* and *SOD3* [Reviewed in (105)], we observed no increase in *SOD2* mRNA levels following HGD (**Figure 5C**). Further work is needed to identify a potential transcription factor responsible for the upregulation of *SOD1* but not *SOD2* mRNA following acute HGD exposure in HBMECs. Finally, and potentially as a consequence of both upregulated VCAM-1 protein and *SOD1* mRNA expression, we observed a significant increase in extracellular H_2_O_2_ following 3h HGD exposure (**Figure 5D**). Similarly, this increase in H_2_O_2_ concentration was not changed with ozanimod treatment which mirrors our findings at both the level of VCAM-1 and *SOD1*. Moreover, it is well known and accepted that H_2_O_2_ due to its small non-polar nature and relatively long half-life can diffuse across biological membranes serves a pleotropic role in cellular function as a second messenger (106) as well as an activator of MMP-9. These findings partially reveal how HGD exposure might be mediating an increase in MMP-9 activity but does not sufficiently elucidate the attenuative role of ozanimod.

### Inhibition of MMP-9 activity and exportation into the extracellular domain

Similar to our investigation into a potential mechanism underlying MMP-9 activation, we next interrogated inhibitory mechanisms of MMP-9 in hopes of identifying a means by which ozanimod exerts its attenuative effects following HGD exposure. We demonstrated that mRNA levels of *RECK* and *TIMP1* decreased following HGD; however, ozanimod did not alter their expression (**Figure 6**). These endpoints were chosen to supplement the positive regulatory mechanisms identified in **Figure 5** demonstrating that in addition to increased H_2_O_2_ production, HGD mediated a downregulation in negative regulatory mechanisms associated with MMP-9. (69, 107–111) A previous study illustrated that RECK protein expression supported neuroprotection and reduced cell death in a mouse model of ischemic stroke (112) however, its neuro-focused viewpoint does not shed light on the impact of the cerebrovasculature. To this end and in support of our findings, an additional in vitro study conducted with human embryonic kidney epithelial cells demonstrated that beginning at 4h of hypoxic conditions, *RECK* mRNA levels were decreased compared to normoxic controls. (113) While RECK has been implicated in the structural inhibition and secretory regulation of MMP-9 (70), it is generally accepted that TIMP1 forms a complex with MMP-9 which prevents its enzymatic activity. (71) In contrast to our findings in **Figure 6B**, a study conducted in a murine model of 0.5h focal cerebral ischemia followed by reperfusion at various times found that the relative preischemic hemisphere concentration of TIMP1 was extremely low and was gradually upregulated beginning at 3h post-reperfusion. (114) However, in the same study the investigators demonstrated an increase in Evan’s Blue extravasation in *TIMP1* ablated mice at 24h post-reperfusion, thus demonstrating its necessity in maintaining BBB integrity. To the best of our knowledge this is the first report to demonstrate a baseline level of *TIMP1* mRNA under normoxic conditions that is acutely decreased following HGD exposure in primary HBMECs; however further investigation is required to fully elucidate the mechanism behind this observation. These data indicate that in addition to increased positive regulation of MMP-9, HGD induces a decrease in the negative regulation of the proteolytic enzyme; however, the regulatory mechanism behind ozanimod remains unclear.

### MMP-9 Membrane Docking

As previously mentioned, the transmembrane glycoprotein CD44 exhibits many different cellular functions; however, in this study we viewed the expression of CD44 through the lens of MMP-9 and endothelial barrier integrity regulation. We hypothesized that ozanimod may be working in part through a CD44 mechanism to attenuate MMP-9 activity and improve barrier integrity, as it has been previously shown that silencing of S1PR1 prevented CD44 and hyaluronan mediated barrier enhancement. (115) The relationship between CD44 and MMP-9 was firmly established by a seminal study which illustrated that docking of proteolytic active MMP-9 to surface CD44 enhanced tumor invasion and growth as well as angiogenesis through a TGF-β mechanism. (116) This gave rise to a multitude of additional studies which have primarily focused on the role of CD44 in tumor microenvironments [Reviewed in (117)] which more recently has shifted to be in the vascular spotlight. [Reviewed in (118)] Counter to our observations of decreased CD44 labeling following 3h HGD exposure (**Figure 7B**), a study in renal microvascular endothelial cells demonstrated an exponential increase in CD44 expression at 1, 3, 7, and 14 days following a 45 min ischemia reperfusion injury induced by bilateral renal artery ligation. (119) Similarly, a model of permanent middle cerebral artery occlusion performed in rats found that both mRNA and protein levels of CD44 in the ipsilateral hemisphere of injury increased beginning at 6h post injury and was maximized at 24h. (120) Additionally, these findings were recently functionally substantiated in a study with iPSC-derived brain microvascular endothelial cells which demonstrated that prolonged oxygen-glucose deprivation exposure for 24h increased CD44 expression and that binding of hyaluronan induced a partial maintenance of endothelial barrier function compared to oxygen-glucose deprivation alone. (121) While these studies do not coincide with our findings here in this study, to the best of our knowledge this is the first report to characterize the acute expression of CD44 in primary HBMECs following HGD exposure. These data could suggest that CD44 undergoes a biphasic response in which expression is acutely decreased and gradually increases following ischemic-like injury.

In addition to measuring levels of labeled CD44, the immunocytochemical staining of this protein sheds light on its nuclear and non-nuclear overlap. Due to the transmembrane nature of CD44, this protein is described as having extracellular, transmembrane, and intracellular domains. [Reviewed in (75)] Soluble release of CD44 has been well accepted as a consequence of proteolytic cleavage of the extracellular domain; however, further investigation into the regulation of CD44 has revealed that the intracellular domain of CD44 is able to translocate to the nucleus and act as a transcription factor following cleavage by membrane type 1-matrix metalloproteinase. (76, 122) Moreover, in a separate study in human umbilical vein endothelial cells, induction of endothelial migration and morphogenic differentiation via type 1-matrix metalloproteinase was shown to be in part dependent on Gi proteins downstream of S1PR1 following sphingosine-1-phosphate stimulation. (123) Suggesting that mechanisms downstream of S1PR1 may also be consequently involved with the regulation of nuclear CD44 expression via membrane type 1-matrix metalloproteinase. The observations made in this study indicate that ozanimod may be attenuating CD44 expression in the nucleus of HBMECs; however, these observations were abolished following 3h HGD exposure (**Figure 7C**). Thus, suggesting that HGD exposure may be differentially regulating various intracellular signaling pathways that potentially negate the effects of ozanimod stimulated S1PR1 signaling with relation to CD44 cleavage which merits further investigation.

### PECAM-1

In a study by Kato et al (16), it was determined that MMP-9 induced cleavage of PECAM-1 in murine hepatic endothelium following ischemia reperfusion injury. These findings were expanded upon a year later which showcased that, deficiencies in MMP-2 expression resulted in upregulated MMP-9 activity and increased transmigration of leukocytes, liver damage, and decreased survival following hepatic ischemic reperfusion injury. (124) Intriguingly, these authors found a significant colocalization of MMP-2 with PECAM-1 in naïve livers which suggests a potential regulatory role of MMP-2 on the stability of PECAM-1 that is disrupted by MMP-9. These studies complement our data which revealed a significant decrease in PECAM-1 staining following 3h HGD exposure which was then attenuated by ozanimod (**Figures 4A & B**), mirroring the activity of MMP-9. Although our data support the hypothesis of an MMP-9 dependent degradation of PECAM-1 following ischemia reperfusion injury, another study conducted in primary rat retinal endothelial cells under hyperglycemic conditions found that MMP-2 mediated cleavage of PECAM-1. (125) While these interpretations contradict the hypothesis of MMP-9 induced cleavage of PECAM-1, the authors found no difference in plasma MMP-9 activity under diabetic conditions, but rather a significant increase MMP-2 activity. Moreover, the diabetic conditions of the study, in which glucose is oversupplied, is almost opposite in nature to the ischemic-like injury model utilized in this study, making a direct comparison between the two outcomes difficult to justify. Additionally, although the authors found multiple cleavage sites on rat PECAM-1 suitable for MMP-2, we utilized the same in silico analysis via PROSPER (126) in this study to demonstrate that there are significantly more sites on human PECAM-1 that can be cleaved by MMP-9 compared to MMP-2 and MMP-3 (**Supplemental Figure 1**). Due to the overlapping substrate material degraded by both MMP-9 and MMP-2 it is possible that human PECAM-1 is degraded by MMP-2; however, in this study we did not observe an increase in MMP-2 activity and surmise that MMP-9 is cleaving PECAM-1 and ozanimod may be attenuating this response via blockade of MMP-9 activity.

In addition to being a marker of MMP-9 activity in this study, we utilized PECAM-1 levels as an indicator of endothelial barrier function. It has been previously observed that the lack of PECAM-1 in primary mouse brain microvascular endothelial cells resulted in an impaired endothelial barrier as measured by trans-endothelial electrical resistance (TEER). (10) Moreover, gerbil brain microvascular endothelial cells exposed to 6h of oxygen glucose deprivation and 12h of reperfusion, showed reduced levels of PECAM-1 as measured by both immunocytochemical and western blot analyses. (127) These previous findings align with the observations in our study which demonstrate that 3h HGD exposure reduced levels of PECAM-1 (**Figure 4B**). Additionally, our western blot observations also shed light on the role of PECAM-1 in forming homophilic interactions which was first hypothesized by Albelda et al (128) and later confirmed by a series of studies to be necessary for its concentration at intercellular junctions. (129–131) This subcellular localization in addition to the relative amount of PECAM-1 has been postulated to regulate endothelial barrier integrity. (132) We hypothesized that the subpopulation of PECAM-1/PECAM-1 detected in our studies at ∼240-260kDa corresponded to the dimerized PECAM-1 formed from homophilic interactions between neighboring endothelial cells. In concordance with the previous observations, we previously observed an HGD-mediated decrease in HBMEC TEER (40) and a corresponding decrease in PECAM-1 (**Figure 4**), suggesting that the maintenance of PECAM-1 via ozanimod is in part responsible for preserving endothelial barrier integrity following ischemic-like injury potentially through the attenuation of MMP-9 activity. Overall, these findings prompted a two-pronged interpretation. 1) The observed decrease in CD44 correlated with a decreased confinement of MMP-9 which 2) could have resulted in an increased ability to degrade structures integral to endothelial barrier function.

### Claudin-5 and ZO-1

To test the latter half of our conclusion we assessed levels of the integral endothelial barrier protein, claudin-5. We chose this protein as it has been well established as being highly expressed in brain endothelial cells at the BBB and shown to be necessary for maintenance of BBB integrity. (8, 133) Moreover, as previously conducted with PECAM-1 (**Supplemental Figure 1**), an in-silico prediction of claudin-5 protease cleavage sites was conducted via PROSPER which revealed that MMP-9 is capable of mediating degradation of both isoforms of claudin-5 protein (**Supplemental Figure 2**). In line with our hypothesis and mirroring our findings of PECAM-1, we observed a notable decrease in claudin-5 protein levels following HGD exposure that was then attenuated by ozanimod, potentially mediated by attenuation of MMP-9 (**Figure 8A**). These results are substantiated by findings in whole-cell extracts from rat brains that underwent a 1.5h middle cerebral artery occlusion followed by 3h of reperfusion, which illustrated a decrease in claudin-5 protein levels compared to sham animals and that administration of a broad-spectrum MMP inhibitor reversed this response. (134) Moreover, in a separate study at 6h post reperfusion following a 2h middle cerebral artery occlusion within a rat model of ischemic stroke, a decrease in both claudin-5 and ZO-1 proteins was observed and functionally correlated with both increased Evan’s Blue extravasation and brain water content. (135) These results correspond with the significant decrease in ZO-1 protein levels detected in this study following HGD exposure (**Figure 8B**). However, unlike claudin-5, the HGD-mediated decrease in ZO-1 levels was not attenuated by ozanimod which further supports our hypothesis that ozanimod may be acting to decrease MMP-9 activity. While our findings support the aforementioned hypothesis, it is also possible that ozanimod induces an S1PR1 mediated intracellular signaling cascade that stabilizes the claudin-5 protein during ischemic-like injury which merits further investigation. Collectively these data suggest that ozanimod, potentially through decreased MMP-9 activity (**Figure 3B**) leading to an attenuation of decreased claudin-5 protein levels (**Figure 8A**), lies at the basis of the previously observed beneficial effects on endothelial barrier function. (40)

In summary, our results suggest that HGD exposure induces a concomitant increase in MMP-9 activity and alteration of proteins associated with endothelial barrier integrity in HBMECs and that ozanimod attenuates these HGD-induced responses, thus adding to its therapeutic potential in cerebrovascular protection during the acute phase of ischemic stroke. We acknowledge that the complex pathophysiological cascade shown to occur during AIS cannot be exactly modeled in an in vitro setting and we acknowledge that HBMECs in culture conditions do not exactly mimic the tubular characteristics similar to those found in vivo. However, in vitro studies using human primary cells do allow the investigation of specific basic cellular and molecular mechanisms under conditions of hypoxia or oxygen and glucose deprivation which is similar to what is observed during AIS. Based on our results in this study we posit that ozanimod is in part working at the level of MMP-9 to attenuate HGD-induced decreases in HBMEC barrier integrity. However, due to the complex and interconnectivity of GPCRs, it is also likely that ozanimod elicits multiple responses in primary human brain microvascular endothelial cells to attenuate ischemic stroke-like associated pathologies related to endothelial barrier function. Thusly, future studies are planned to investigate receptor dependence and the specific mechanism(s) of action in the cerebrovasculature including modulation of Akt, PI3K, Rho, and upstream mediators as well as barrier proteins in freshly dissected, uncultured, endothelial cells. Further understanding the role S1PRs play in maintaining the integrity of the cerebrovasculature following an ischemic injury may provide a stronger basis for the selective clinical targeting of S1PR1 with compounds such as ozanimod for AIS.

## ACKNOWEDGEMENTS

Supported by American Heart Association 19AIREA34480018 (R.J. Gonzales), Valley Research Partnership Award VRP37 P2 (R.J. Gonzales), and Valley Research Partnership Award VRP55 P1a (T.S. Wendt and R.J. Gonzales).

---

**Supplemental Figure 1.**
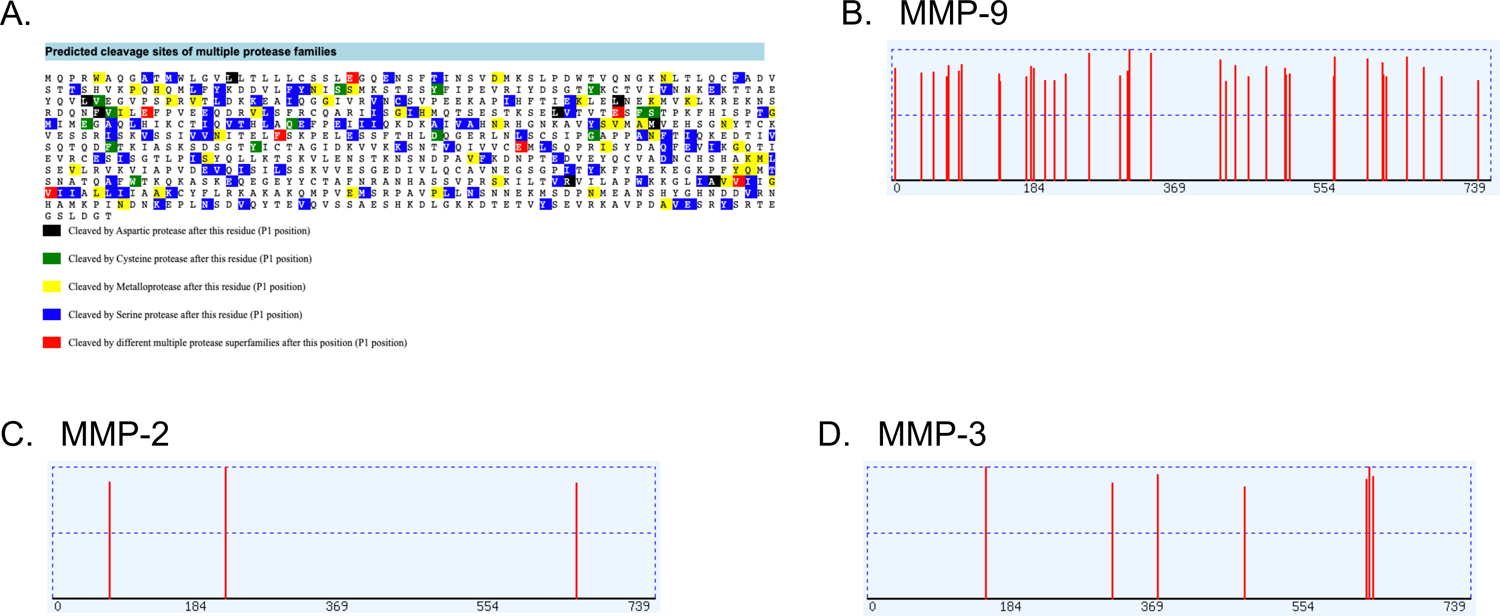
In silico analysis of cleavage sites within human PECAM-1 via PROSPER. (A) Whole sequence of human platelet endothelial cell adhesion molecule 1 (PECAM-1) highlighted at specific amino acid sites to indicate potential cleavage by aspartic proteases (black), cysteine proteases (green), metalloproteases (yellow), serine proteases (blue), and different multiple protease superfamilies (red). (B-D) Within the PECAM-1 sequence highlighted yellow, specific sites with a cleavage probability score greater than 0.8 are shown as red lines and represent independent, potential sites of human PECAM-1 cleavage by MMP-9 (39 independent sites) (B), MMP-2 (3 independent sites) (C), and MMP-3 (7 independent sites) (D).

**Supplemental Figure 2.**
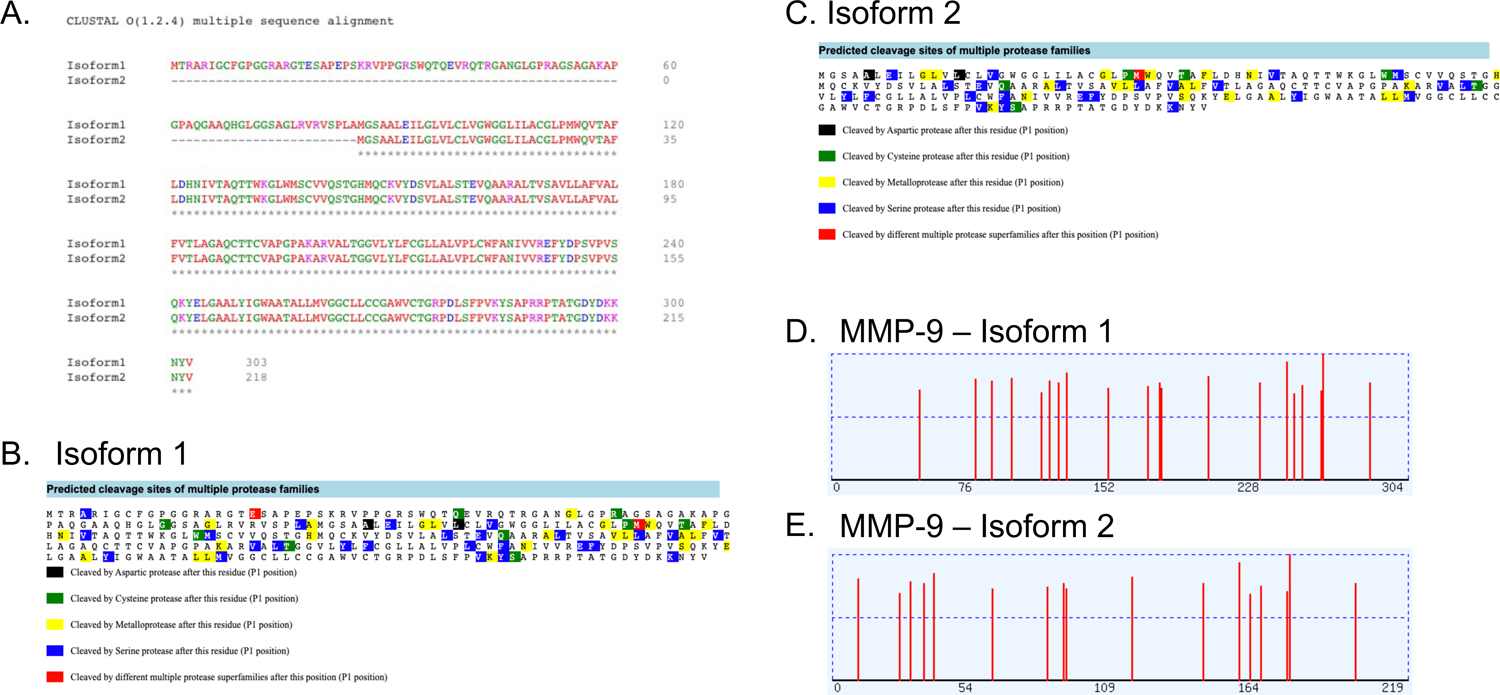
In silico alignment of claudin-5 isoforms and analysis of cleavage sites. (A) Whole proteomic sequence alignment of claudin-5 isoforms 1 and 2 demonstrating total length of isoform 1 (303 amino acids) and isoform 2 (218 amino acids). Asterisks indicate overlapping amino acids between the two claudin-5 isoforms. (B) Whole sequence of human claudin-5 isoform 1 highlighted at specific amino acid sites to indicate potential cleavage by aspartic proteases (black), cysteine proteases (green), metalloproteases (yellow), serine proteases (blue), and different multiple protease superfamilies (red). (C) Whole sequence of human claudin-5 isoform 2 highlighted at specific amino acid sites to indicate potential cleavage by aspartic proteases (black), cysteine proteases (green), metalloproteases (yellow), serine proteases (blue), and different multiple protease superfamilies (red). (D-E) Within the claudin-5 isoform 1 and 2 sequences highlighted yellow, specific sites with a cleavage probability score greater than 0.8 are shown as red lines and represent independent, potential sites of human claudin-5 isoform 1 cleavage by MMP-9 (20 independent sites) (D) and claudin-5 isoform 2 cleavage by MMP-9 (17 independent sites) (E).

